# Robust causal gene network estimation for large-scale single-cell perturbation screens using reduced control function

**DOI:** 10.64898/2026.04.20.719759

**Authors:** Changhao Ge, Hongzhe Li

## Abstract

Single-cell CRISPR perturbation screens provide a foundation for causal discovery in gene regulatory networks, but existing methods struggle with latent confounding, count-valued expression data, and high-multiplicity-of-infection (high-MOI) designs. We introduce RICE, a unified framework that addresses all three challenges. At its core is a reformulated control function tailored to the network-estimation setting, which restores robustness to exclusion-restriction violations that are pervasive in real CRISPR screens and that cause standard control function approaches to fail. RICE pairs this estimator with a constrained negative binomial model and a differentiable acyclicity penalty, accommodating hard and soft interventions and natively supporting high-MOI designs within a single, GPU-scalable model. Across extensive synthetic benchmarks, RICE consistently outperforms existing methods and remains stable under strong confounding, exclusion violations, and high-MOI conditions. Applied to CRISPRi screens, RICE achieves stronger causal-discovery performance on held-out data. RICE further reconstructs the canonical interferon (IFN)-*γ* signaling pathway in melanoma cells upon immune stimulation, and nominates stable regulatory candidates that conventional significance tests may overlook due to limited statistical power. Together, these results establish RICE as a robust and scalable framework for causal discovery in single-cell perturbation genomics.

## INTRODUCTION

Single-cell perturbation screens have become a central tool for studying gene regulation, combining pooled CRISPR perturbations with single-cell transcriptomic readouts to measure how individual cells respond to targeted genetic intervention^1^. Unlike observational single-cell data, these experiments generate interventional variation, making them a natural foundation for causal discovery^2–4^. The central goal in this setting is to reconstruct causal gene networks — directed graphs in which edges encode regulatory influence rather than undirected association — typically represented as directed acyclic graphs (DAGs). Recovering such structure is essential for distinguishing direct regulatory effects from those mediated through downstream pathways, and for understanding how perturbations propagate through regulatory programs^5–7^.

A persistent obstacle to this goal is unmeasured confounding. Latent factors such as technical artifacts, cell-state heterogeneity, and other unobserved variables can influence many genes simultaneously, inducing dependencies that are statistical but not causal^8^. If these spurious dependencies are not explicitly modeled, they can be misinterpreted as regulatory edges, and the inferred network is corrupted by false positives. This problem is intrinsic to single-cell data, and any method intended for real perturbation screens must contend with it.

Control function methods, well established in econometrics for estimation under endogeneity^9^, offer a principled route to addressing latent confounding. They exploit the fact that CRISPR perturbations, being assigned independently of unmeasured factors, can serve as valid instruments. However, control functions were developed for equation-level estimation — recovering a single causal relationship — and their direct extension to network estimation does not behave well. The full control function regresses each gene on the entire perturbation vector, so any exclusion-restriction violation in a single perturbation contaminates the fitted values for every downstream gene and propagates into the corresponding edge estimates. Because such violations are pervasive in real CRISPR screens, the natural tool for the confounding problem fails in precisely the DAG-estimation setting.

Here we show that a *reduced* control function resolves this failure. By residualizing each gene only on its own perturbation indicator, rather than on the full set of perturbations, the reduced formulation isolates the directed causal influence among genes while still absorbing latent confounding. This modification trades a modest amount of statistical power, readily afforded by the large sample sizes typical of single-cell screens, for a substantial gain in robustness, making confounding-robust causal discovery tractable at large DAG scale. We develop this idea into RICE (Robust Identification of Causal Entanglements), an integrated framework for causal gene network estimation in large-scale single-cell perturbation screens.

RICE naturally accommodates the conditions that modern screens actually present. It models count-valued transcript data directly through a constrained negative binomial GLM, avoiding both the linearity assumptions of recent interventional methods such as inspre^10^, IBCD^11^, and dotears^12^, and the restrictive distributional assumptions of count-based approaches such as ODS^13^ and ZIPBNs^14^. It handles both hard interventions, such as CRISPR knockout, which removes upstream regulation of the targeted gene, and soft interventions, such as CRISPRi and CRISPRa, which shift the conditional distribution of the targeted gene while preserving its upstream dependence^15–18^ (Figure 1B); this distinction destabilizes methods tailored to only one regime, whereas RICE captures both within a single parameterization. RICE also supports high-multiplicity-of-infection (high-MOI) designs natively. Low-MOI screens, in which each cell receives at most one perturbation, scale linearly with library size and are efficient for small panels but prohibitively expensive at genome scale; high-MOI designs deliver multiple perturbations per cell and have emerged as a more scalable alternative^6^ (Figure 1C). Many computational pipelines analyze one perturbation at a time, ignoring the combined regulatory effects of additional perturbations present in the same cell^19–21^; RICE instead models the joint effect of cooccurring perturbations directly. This joint treatment respects the biological structure of high-MOI experiments and translates into improved performance in these settings.

**Figure 1:**
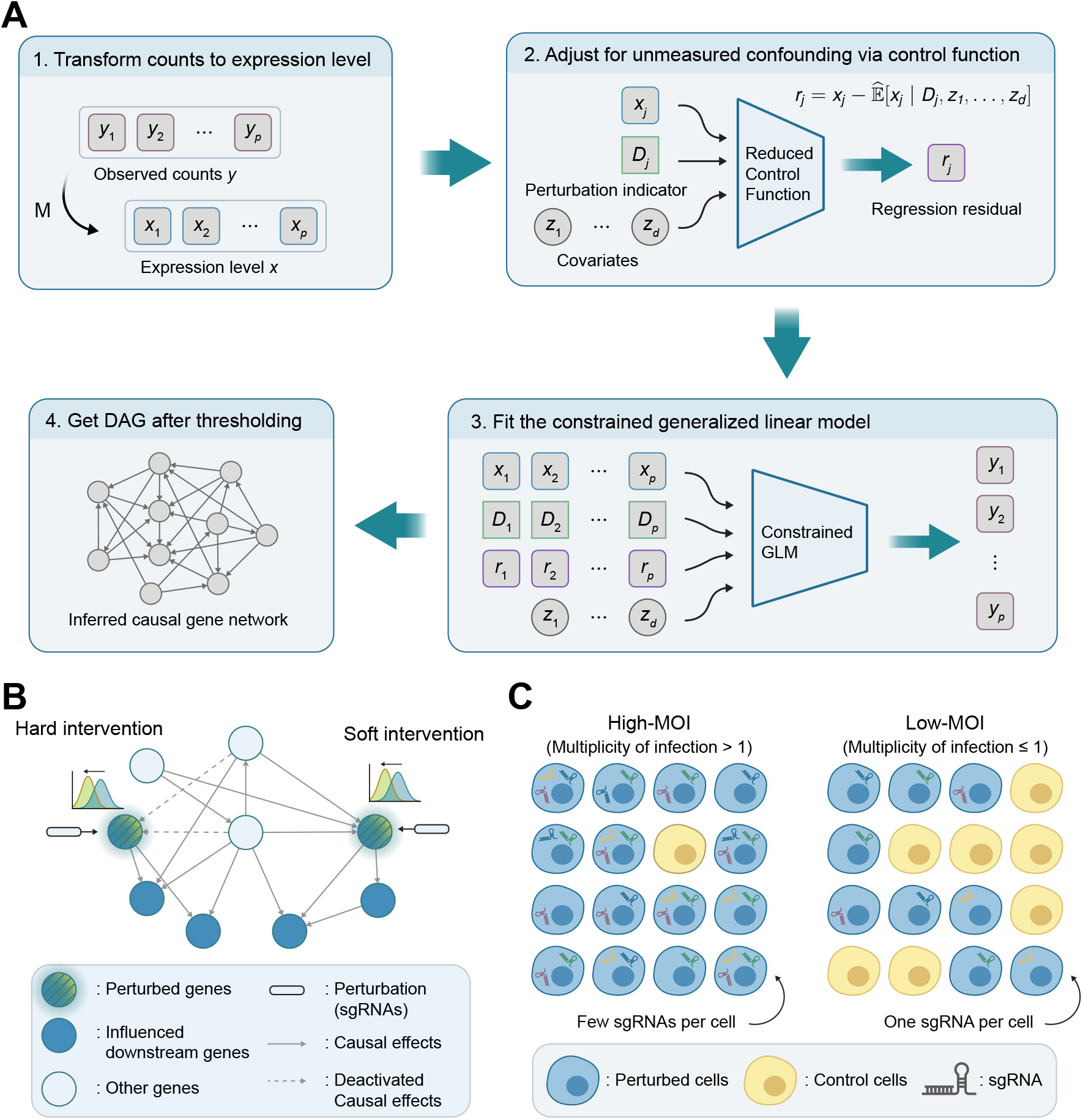
Overview of the RICE framework and experimental paradigms. **(A)** Schematic of the RICE pipeline. For each gene, expression levels are regressed against perturbation indicators and observed confounders to derive residuals. The resulting data are modeled using a generalized linear model (GLM) subject to a DAG constraint. **(B)** Schematic of causal gene networks under soft and hard interventions. Soft interventions preserve the underlying network topology, whereas hard interventions (e.g., gene knockout) eliminate all incoming edges to the targeted gene. **(C)** Illustration of low- and high-multiplicity of infection (MOI) screens. In low-MOI regimes, perturbed cells receive a single perturbation, whereas high-MOI screens allow multiple concurrent perturbations per cell.

## RESULTS

### Model overview

RICE estimates a causal gene network in four steps (Figure 1A), built around a generalized linear model (GLM) framework. Suppose we observe read counts ***y*** = (*y*_*j*_) together with covariate ***z***. Because modeling the relationships among raw counts directly is numerically unstable, RICE first transforms the counts into continuous expression levels ***x*** = (*x*_*j*_) via a non-decreasing univariate mapping. This mapping is applied only to the GLM predictors; the response still uses the raw counts ***y***. RICE then constructs a reduced control function to absorb unmeasured confounding, fits a constrained negative binomial GLM that enforces a DAG structure on the inferred causal matrix, and thresholds the learned weights to obtain the final network.

The reduced control function step is the conceptual core, which residualizes each gene on its own perturbation. Concretely, the expression of each gene *j* depends on its parents, on its own perturbation (sgRNAs), and on unmeasured confounders ***z***_um_. The reduced control function is defined, for each gene *j*, as

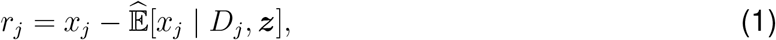

where the conditioning set includes the gene’s own perturbation indicator *D*_*j*_ and observed covariates ***z***. Because *D*_*j*_ is assigned independently of ***z***_um_ and is exogenous with respect to every upstream gene, *E* [*x*_*j*_ | *D*_*j*_, ***z***] isolates the variation in *x*_*j*_ attributable to perturbing *j*. The residual *r*_*j*_ therefore carries no perturbation-induced regulatory signal and reflects only latent confounding and intrinsic noise. Conditioning the second-stage model on ***r*** = (*r*_*j*_) consequently yields regression coefficients that serve as proxies for the causal effects and thus preserves the causal target.

Under this formulation, the conditional mean of gene *j* is modeled as

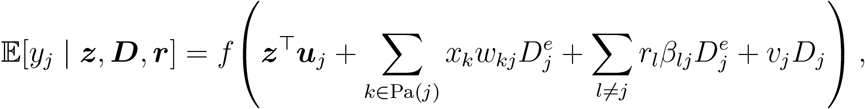

where *f* is the GLM link function, ***u***_*j*_ is the coefficient of covariates for gene *j*, Pa(*j*) denotes the parent set of gene *j, w*_*kj*_ is the direct causal effect of gene *k* on gene *j, β*_*lj*_ is the coefficient of the control function residual from gene *l, v*_*j*_ is the effect of perturbing gene *j*, and *D*^*e*^ encodes the intervention type 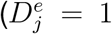 for soft interventions and 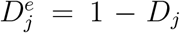 for hard interventions,which remove upstream regulation of the perturbed gene). The weight *w*_*kj*_ measures the direct effect of gene *k* on gene *j* after conditioning on the residuals. The cost is a modest reduction in statistical power, since the first stage introduces additional variables; however, as we show later (Figure 4A), we find it does not materially affect recovery of strong causal edges.

RICE then fits a regularized negative binomial GLM using the full set of variables, including the residuals ***r*** = (*r*_*j*_), while constraining the inferred causal matrix to a DAG. The negative binomial likelihood accommodates the overdispersion characteristic of single-cell count data, and the same parameterization handles both hard and soft interventions, with only the definition of ***D***^*e*^ differing between regimes; extensions to other count-based distributions are described in Note S1. Because the combinatorial DAG constraint is intractable, RICE replaces it with a differentiable acyclicity constraint^22^, making the objective smooth, gradient-optimizable, and GPU-accelerable. These properties are essential for scaling to the hundreds of genes and hundreds of thousands of cells in modern screens. After optimization, RICE applies change-point detection to the sorted edge weights to separate true causal signal from a tail of near-zero background weights, yielding the final DAG (see Methods).

### The reduced control function is preferable for Perturb-seq analysis

The formulation in (1) is termed the *reduced* control function to distinguish it from its full counterpart, in which the residual conditions on all perturbations,

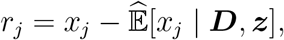

recovering the standard control function. Both formulations are valid under the standard assumptions of the instrumental variable framework. Under the conditions typical of modern single-cell perturbation screens, however, the two exhibit different properties, and we find the reduced control function preferable. The principal reason is its greater robustness to model misspecification in practice, a regime in which the full control function breaks down.

The instrumental variable framework requires that each perturbation *D*_*j*_ affects downstream gene expression only through the targeted gene *x*_*j*_, an idealization rarely met in practice. Real CRISPR perturbations can produce effects on non-target genes through pathways that bypass the targeted gene, including off-target sgRNA activity, perturbation-induced changes in cellular state such as DNA damage responses, and effects mediated by unmeasured intermediaries — proteins, chromatin states, and non-coding RNAs — that fall outside the gene expression panel^23,24^. When such violations occur, *D*_*j*_ retains predictive power for genes other than *x*_*j*_ along paths that do not pass through *x*_*j*_.

The full control function, which regresses each gene on the entire perturbation vector, propagates these out-of-target signals into the first-stage fitted values 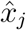 used for downstream genes, contaminating the second-stage edge estimates. The reduced control function avoids this pathway: because 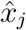 depends only on *D*_*j*_, out-of-target effects of other perturbations cannot enter 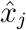 and therefore cannot propagate into the edge estimates incident to gene *j*.

We emphasize that, under such violations, neither formulation recovers the structural causal graph in the strict sense. The reduced formulation nevertheless yields edge estimates that remain locally interpretable and empirically stable, whereas the full formulation produces estimates that depend non-locally on the entire perturbation design and can be substantially biased.

We confirm this property empirically through simulations with such violations (Figure 2), evaluating performance using positive predictive value (PPV), true positive rate (TPR), Matthews correlation coefficient (MCC), and structural Hamming distance (SHD); full simulation details are provided in Methods. In the presence of off-target effects, the full control function exhibits substantial degradation in edge recovery, whereas the reduced formulation remains stable. The deterioration is driven primarily by an excess of false positives under the full formulation, which depresses PPV from over 0.8 to roughly 0.65; combined with a marginal decrease in TPR, this loss propagates to MCC and SHD. The same pattern emerges in our real-data analyses (Figure S1). Because exclusion violations of this form are difficult to rule out in practice, this robustness is a substantive advantage in Perturb-seq settings.

**Figure 2:**
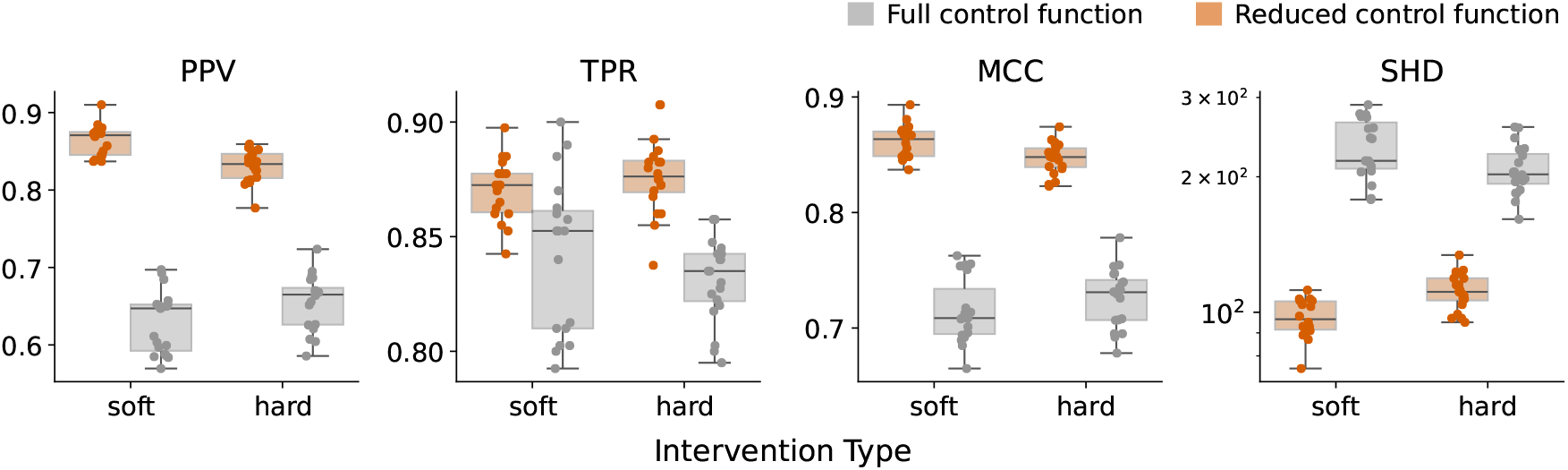
The reduced control function yields more stable performance than the full control function under exclusion-restriction violations. Each point represents a single simulation replicate with 10,000 cells, 100 genes and 400 true causal edges; boxplots summarize 20 replicates per method. Abbreviations are defined in Methods.

### Accurate and efficient DAG recovery on synthetic data

To evaluate the performance of RICE in causal discovery, we applied it to synthetic Perturb-seq data and compared it with inspre, IBCD and dotears. For a fair comparison, the log(1 + *y*) transformation was applied to inspre, IBCD, and dotears for three reasons: these methods assume continuous data, the transformation reflects the data-generating process, and it outperformed other common choices such as a linear transformation for these methods. The simulation details are reported in Methods.

Figure 3A shows representative examples of three manually curated network topologies: hub, chain, and clusters. We further evaluated the models under three data-generating distributions: Poisson, negative binomial, and Poisson log-normal (Figure 3B). RICE ranked first or nearly first across evaluation metrics in most experimental settings. We attribute this robust performance, even under model misspecification, to the flexibility of the negative binomial distribution (see Methods). Although inspre achieved high TPR, this came at the cost of numerous false discoveries, resulting in substantially lower PPV and MCC. By contrast, IBCD achieved high PPV but had low TPR, suggesting overly conservative behavior. Finally, dotears performed poorly in both PPV and TPR, indicating a limited ability to recover the underlying regulatory structure in this setting. Additional simulation settings, including alternative graph topologies, weak perturbation effects, weak causality, and larger network scales, are summarized in Figure S2.

**Figure 3:**
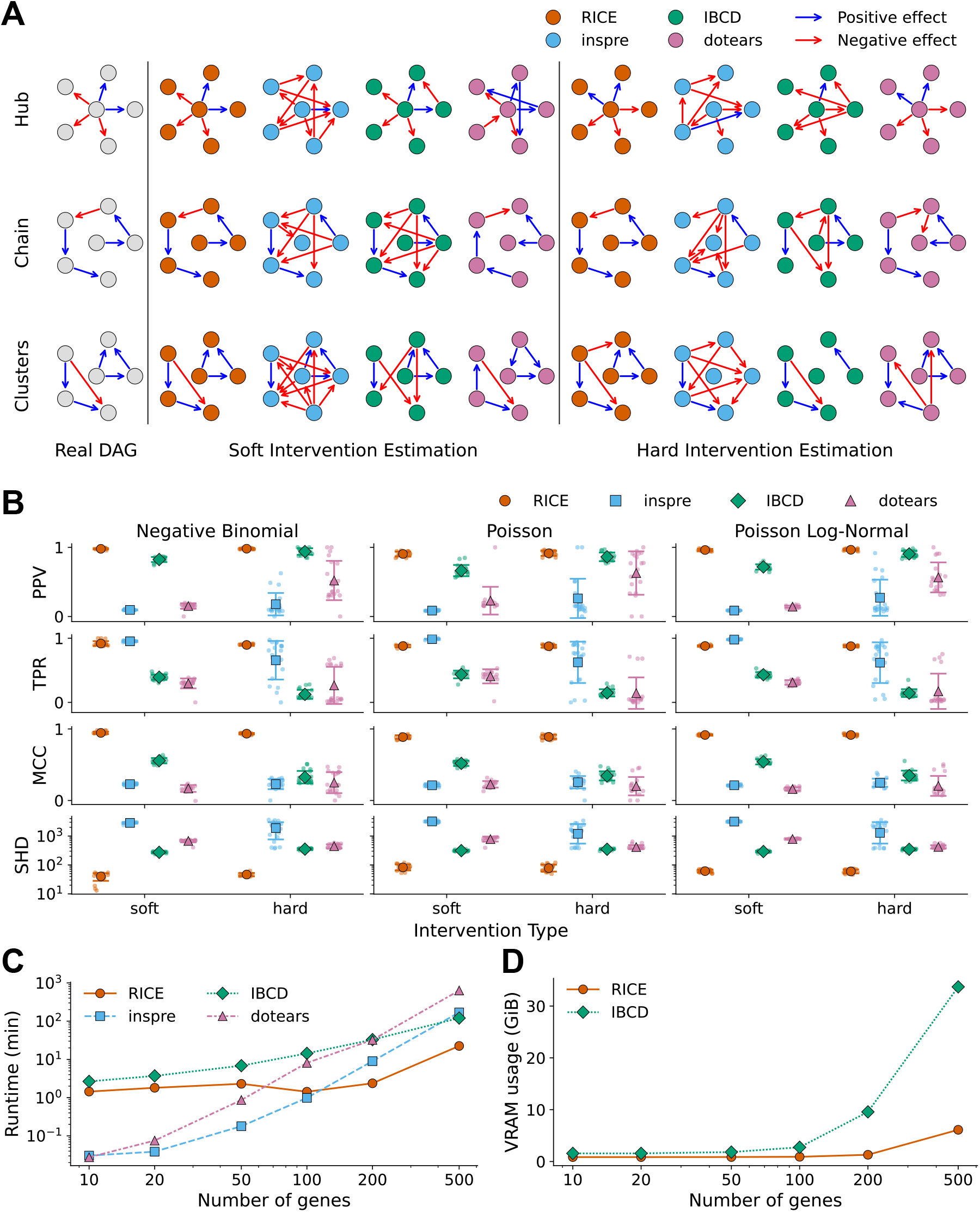
RICE outperforms existing methods in accuracy and efficiency. **(A)** Representative examples of DAG recovery using 500 cells and 6 genes. Three distinct network topologies were curated, with data generated under both soft and hard intervention regimes. **(B)** Benchmarking results on Erdo’’ s–Re’nyi graphs across three data-generating processes. Each point represents a single replicate with 10,000 cells, 100 genes and 400 true causal edges. Error bars denote one standard deviation. **(C)** Computational runtime as a function of the number of genes. **(D)** GPU memory consumption of RICE and IBCD as a function of the number of genes. For (C) and (D), markers represent the mean across 10 replicates. Abbreviations are defined in Methods.

An additional pattern is that inspre and IBCD were less stable under hard interventions, whereas dotears performed worse under soft interventions. This is consistent with the methods’ underlying intervention models: dotears identifies exogenous variance from the edge-breaking structure of hard interventions, whereas inspre and IBCD model perturbations as changes that leave a gene’s incoming edges intact — i.e., soft interventions. Each method is therefore matched to one regime, while RICE accommodates both soft and hard interventions and maintains stable performance across settings.

Another advantage of RICE is its computational efficiency in large-scale settings, driven by GPU-accelerated matrix operations. Figure 3C summarizes algorithm runtime as a function of the number of genes. CPU-based methods such as dotears and inspre were efficient on small networks, but their runtimes increased rapidly with network size. By contrast, RICE showed markedly better scalability, with little noticeable increase in runtime for networks containing up to 200 genes. For networks with 500 genes, CPU-based approaches typically required several hours, whereas RICE completed in approximately twenty minutes.

Although IBCD also leverages GPU computation and showed similar scaling behavior, its reliance on posterior sampling introduced substantial overhead, leading to longer runtimes in practice. In addition, IBCD incurred substantially higher GPU memory consumption (Figure 3D). For example, on datasets containing 100,000 cells and 500 genes, RICE used less than 10 gibibytes (GiB) of GPU memory—within the capacity of many consumer-grade GPUs—whereas IBCD required more than 30 GiB. These results suggest that, under the same hardware constraints, RICE can accommodate substantially larger datasets than IBCD.

### Robustness to unmeasured confounding via reduced control function

A key concern in perturbation-based causal discovery is whether the inferred graph remains reliable in the presence of unobserved factors that jointly affect multiple genes. To examine this, we conducted an additional simulation study in which the number of unmeasured confounders was increased from 2 to 10 under both soft and hard interventions. As shown in Figure 4A, RICE with reduced control function (RCF) remained robust as latent confounding became stronger, with all metrics staying relatively stable across settings. In contrast, competing methods either lost precision or recovered fewer true signals as the effect of unmeasured confounders increased, leading to poor performance under strong confounding.

**Figure 4:**
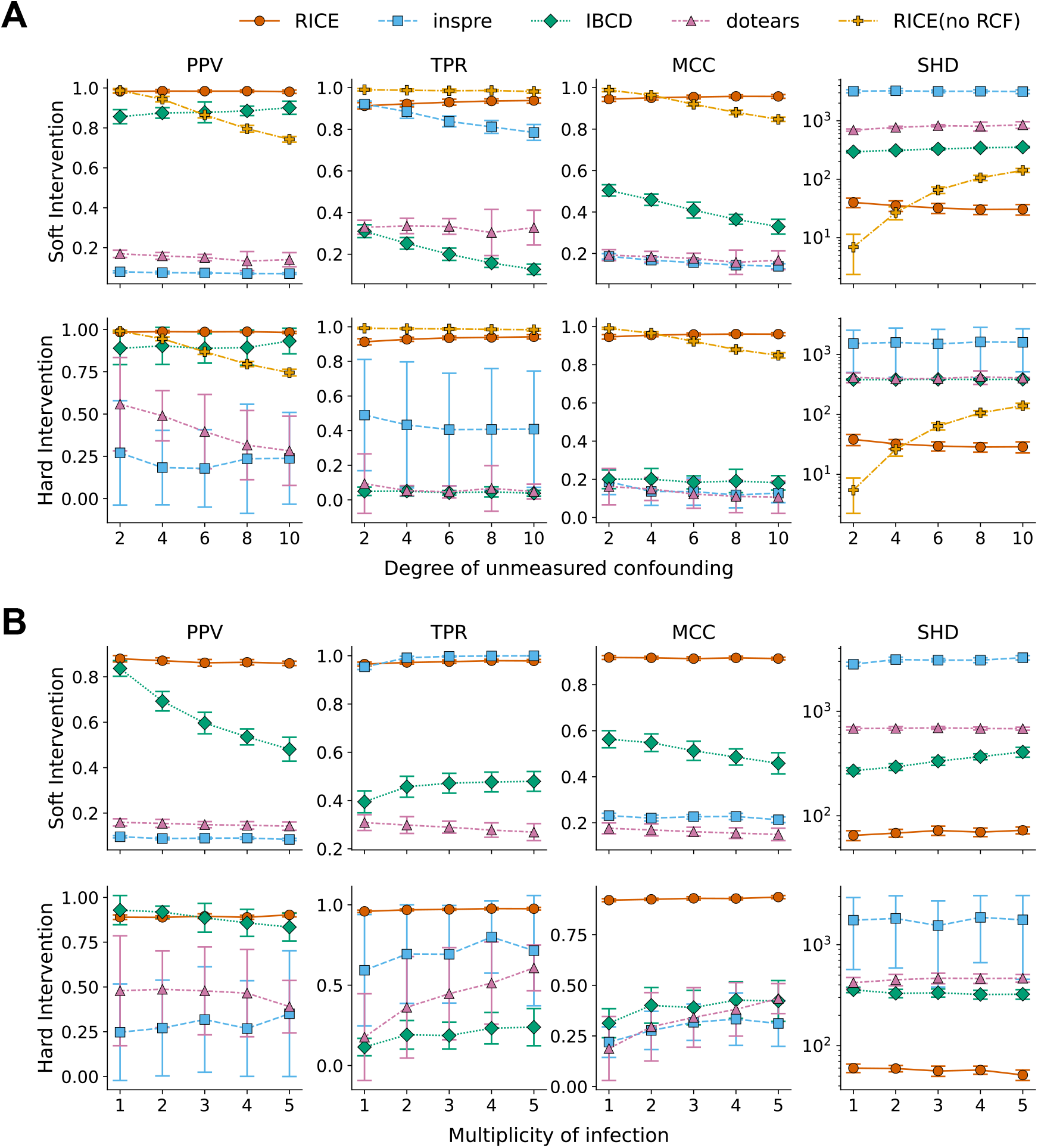
Simulation results highlight the robustness of RICE to unmeasured confounding and high multiplicity of infection (MOI). **(A)** Performance comparison across a range of i.i.d. unmeasured confounders. RCF: reduced control function. **(B)** Evaluation of model performance under increasing MOI. For both panels, each simulation is based on 10,000 cells, 100 genes and 400 true causal edges. Markers represent the mean across 20 simulation replicates, and error bars denote one standard deviation.Abbreviations are defined in Methods.

Figure 4A also serves as an ablation study of RCF. When the number of unmeasured confounders was small, RICE without RCF showed slightly better performance, since it avoids the loss of statistical power introduced by residualization. However, its performance deteriorated rapidly as confounding increased. In particular, removing RCF led to more false positives, indicating that confounding effects were misinterpreted as causal relationships. As a result, although TPR remained consistently high, the model without RCF showed a marked decline in PPV and MCC, together with a substantial increase in SHD. These results highlight a tradeoff between power and robustness and show that RCF is essential for maintaining reliable graph recovery in the presence of strong unmeasured confounding like those in the Perturb-seq data.

### Extension to high-MOI screens

To evaluate whether RICE remains effective beyond the conventional low-MOI regime, we conducted an additional simulation study under increasing MOI, in which individual cells may carry multiple perturbations simultaneously. As described in the model, RICE naturally accommodates this setting through an additive formulation of perturbation effects without altering the overall estimation framework. The competing methods, by contrast, are formulated for single-perturbation data and provide no native mechanism for multiple co-occurring perturbations, so comparing them at high-MOI requires adapting the input. We adopted the most direct and transparent adaptation available: each cell carrying *m* perturbations was expanded into *m* pseudo-observations, one per perturbation, so that every perturbation–response pair is presented to each method exactly as it would appear in low-MOI data. This construction does not explicitly model perturbation co-occurrence, but it supplies each method with the full set of single-perturbation signals it was designed to exploit, rather than discarding high-MOI cells or forcing the methods to fail.

As shown in Figure 4B, RICE remained robust as MOI increased, with performance staying relatively stable across evaluation metrics over a broad range of settings. By contrast, under soft interventions, IBCD showed declining performance with increasing MOI, suggesting that it does not generalize well to high-MOI data. Although the other two competing methods did not exhibit a pronounced MOI-dependent decline, their performance remained consistently poor across all settings, indicating limited effectiveness even before entering the high-MOI regime. Furthermore, performance was continuous from MOI=1 onward, with no jump where cells begin carrying multiple perturbations, indicating that the baselines were not handicapped, and RICE’s advantage is genuine rather than an artifact of a weakened comparator. Notably, under hard interventions, the TPR and MCC of all methods improved marginally as MOI increased. This reflects the fact that simultaneous perturbation of multiple genes removes their dependence on upstream regulators, thereby simplifying recovery of the remaining graph.

Taken together, these results highlight the flexibility of RICE across a broad range of MOI settings. This robustness makes it a practical framework for causal discovery in increasingly complex single-cell perturbation studies.

### Uncovering causal structure in CRISPRi screens of K562 and RPE1 cells

We benchmarked model performance on CRISPRi screens in K562 and RPE1 cells targeting essential genes with single-gene perturbations^19^, treated as soft interventions in RICE. K562 is a human immortalized myelogenous leukemia cell line, whereas RPE1 is a non-cancerous hTERT-immortalized retinal pigment epithelial cell line. After preprocessing, we split the data into two parts: 80% of the cells were used for model training, and the remaining 20% were reserved for benchmarking. The perturbation proportions in each split were preserved.

RICE identified 2,850 edges in K562 (Supplementary Data S1) and 1,191 edges in RPE1 (Supplementary Data S2). For comparison, inspre, IBCD, and dotears identified 3,252, 4,314, and 4,817 edges in K562, and 1,813, 1,250, and 2,348 edges in RPE1, respectively. RICE and IBCD exhibited similar scale-free-like degree distributions (Figures S3 and S4), characterized by a concentration near zero and a small number of highly connected genes. This pattern is consistent with prior reports of power-law behavior in gene regulatory networks^25–27^. RICE produced the shallowest network, with a maximum depth of 12 layers and most genes located near the top, in line with previous findings of shallow and largely feedforward structure of transcriptional networks^28–30^. In contrast, inspre and dotears yielded substantially deeper networks (depth *>* 40).

We first examined whether inferred edge weights were associated with total causal effects in the held-out data. In CRISPRi, gene repression is expected to decrease downstream expression under positive regulation and increase it under negative regulation, implying a negative correlation between edge weights and total effects. This relationship was strongly observed for RICE, with *R*^2^ = 0.71 in K562 and *R*^2^ = 0.64 in RPE1 (Figure 5A), compared to at most *R*^2^ = 0.35 for other methods. Moreover, 95% of RICE-inferred edges had signs opposite to their corresponding total effects, versus less than 86% for competing methods.

**Figure 5:**
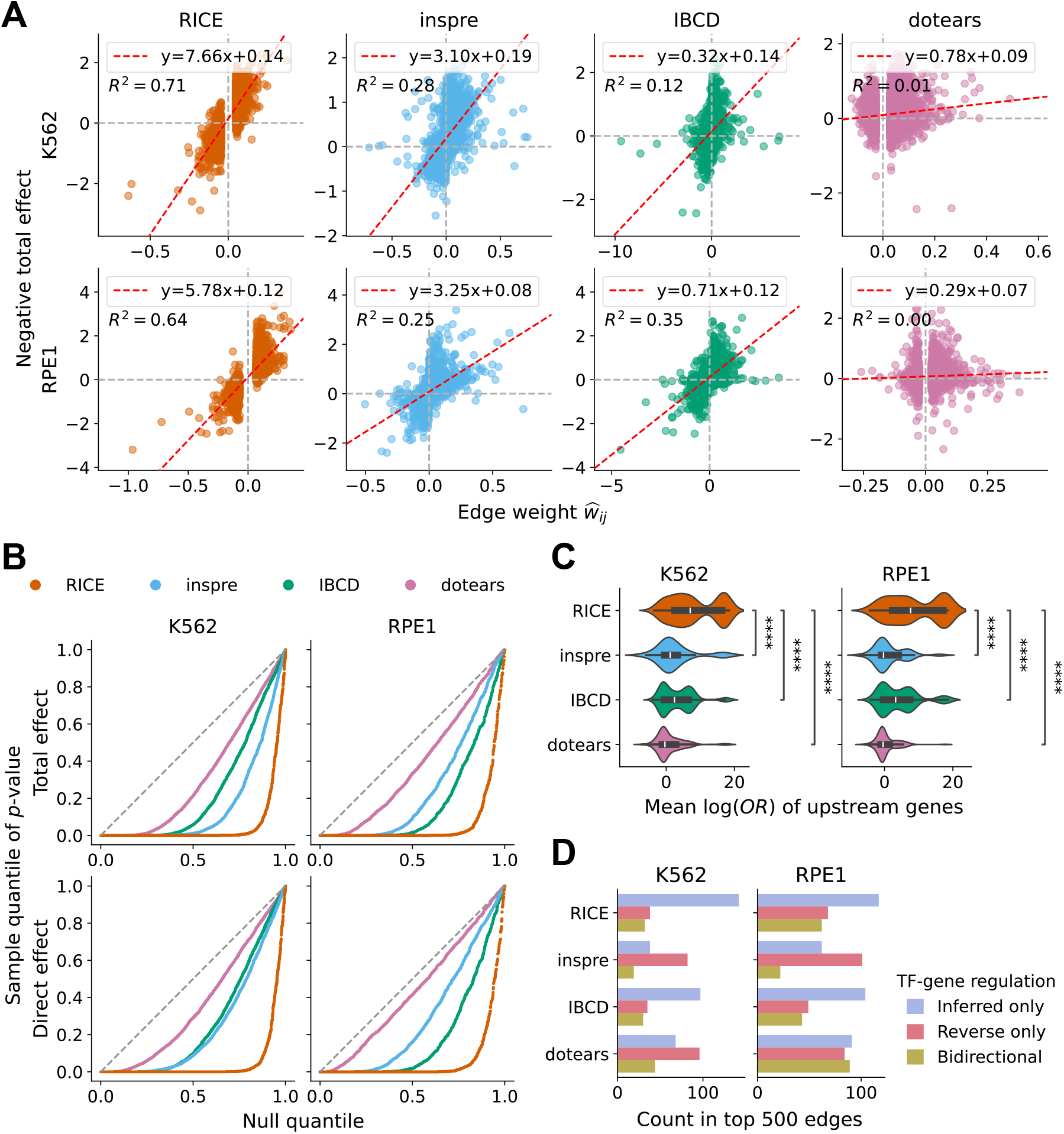
RICE shows superior causal discovery performance on held-out K562 and RPE1 cells. **(A)** Correlation between negative total effects of regulator–target pairs and inferred edge weights. Red dashed lines represent linear regression fits; *R*^2^ denotes the coefficient of determination. **(B)** Quantile–quantile plots comparing *p*-values for total and direct causal effects against the theoretical uniform distribution under the null. **(C)** Mean log odds ratios (OR) for the enrichment of inferred upstream genes that, upon perturbation, induce significant differential expression of the target gene (Mann–Whitney *U* test, *p <* 0.01). ORs were calculated relative to a null distribution of randomly sampled gene sets. Comparisons are based on the Mann–Whitney *U* test. **(D)** Number of transcription factor (TF)–target gene regulatory interactions among the top 500 inferred edges supported by external knowledge bases. Definitions of total effect, direct effect and OR can be found in Methods.

Next, we evaluated inferred edges using both total and direct causal effects (Figure 5B). Because this analysis was performed on held-out data with a relatively limited sample size (minimum of 20 perturbed cells), the resulting *p*-values were not expected to be extremely small. Nevertheless, more than 80% of RICE edges were significant (*t*-test, *p <* 0.05) for both effect types in both cell lines, whereas none of the competing methods achieved comparable significance. Additionally, in RICE, edges with larger inferred magnitudes were more likely to exhibit downstream perturbation effects after data normalization (Figures 6A and S5A), providing a natural basis for prioritizing the identified edges for further investigation.

**Figure 6:**
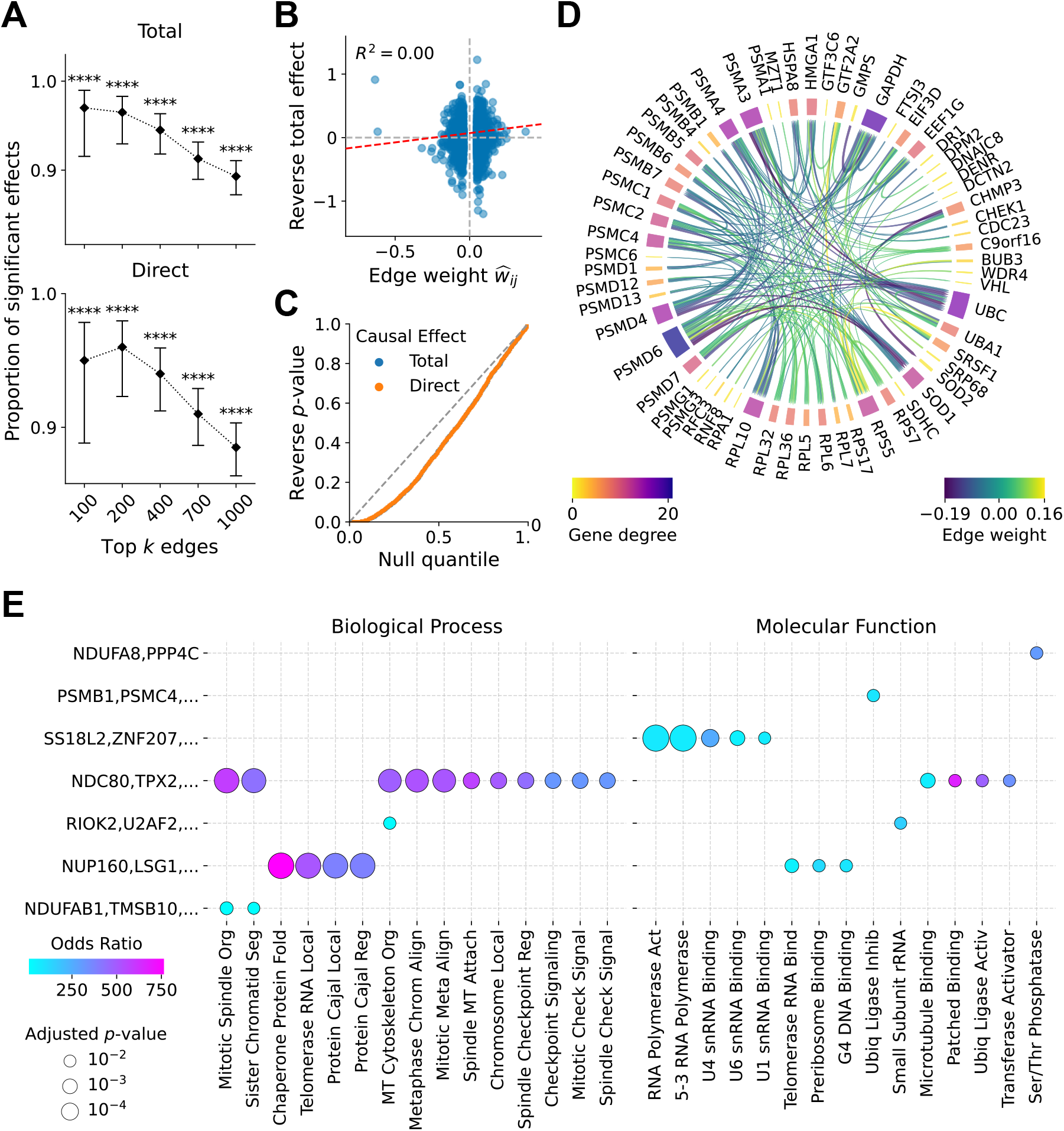
RICE identifies biologically coherent causal structures in K562 cells. **(A)** Proportions of top-ranked edges (from top 1,000 to top 100) exhibiting significant total or direct causal effects (*t*-test, *p <* 0.01). The proportion of validated effects increases with higher edge importance. Error bars denote 95% Wilson binomial confidence intervals. **(B)** Correlation between total causal effects of reversed regulator–target pairs and inferred edge weights. Red dashed lines represent linear regression fits; *R*^2^ denotes the coefficient of determination. **(C)** Quantile– quantile plots of *p*-values for reversed regulator–target pairs compared against the theoretical uniform distribution under the null. **(D)** Representative gene cluster with 61 genes identified via Louvain community detection. **(E)** Gene Ontology enrichment analysis across identified clusters for “Biological Process” and “Molecular Function” categories. The top 15 terms ranked by odds ratio among those with Benjamini–Hochberg adjusted *p <* 0.01 are shown. Only clusters containing at least one enriched term are displayed.

To assess edge directionality, we examined reversed-edge effects by testing whether perturbation of inferred target genes influenced their corresponding upstream regulators. As shown in Figures 6B, 6C, S5B and S5C, such reciprocal effects were not observed for RICE, indicating that the model captures directed causal relationships rather than undirected gene associations. Collectively, these results suggest that RICE effectively recovers both total causal effects and non-mediated interactions, enabling more accurate reconstruction of causal pathways.

For each gene, we analyzed whether its inferred upstream regulators exerted causal effects on its expression; specifically, whether the gene showed expression changes when its identified parent genes were perturbed. To quantify this, we computed the mean log-odds ratio of the fraction of upstream genes with significant differential expression (Mann–Whitney *U* test, *p <* 0.01) (Figure 5C). RICE produced positive values in both datasets (one-sided Wilcoxon signed-rank test, *p <* 10^*−*4^) and significantly outperformed all other methods (Mann–Whitney *U* test, *p <* 10^*−*4^).

Figure 5D summarizes the number of transcription factor (TF)–gene regulatory interactions, together with their inferred directions, supported by TRRUST^31^, PerturbAtlas^32^, OmniPath^33^, and RummaGEO^34^ among the top 500 identified gene pairs for each method. RICE recovered the largest number of annotated interactions in both cell lines while maintaining a relatively low number of incorrectly oriented (reversed) edges, providing additional support for its accuracy in causal discovery.

We further characterized the causal network inferred by RICE in K562 cells, identifying 11 modules via the Louvain algorithm^35^. Module sizes ranged from a two-gene doublet to a 118-gene component (Supplementary Data S1), and several modules captured coherent functional programs with biologically interpretable internal structure. One representative module (Figure 6D), comprising 61 genes, contains a coherent ubiquitin–proteasome substructure: it groups together members of every major proteasome subunit family — PSMA and PSMB (20S core) together with PSMC and PSMD (19S regulatory particle) — alongside the assembly chaperone family PSMG, recovering the structural architecture of the 26S proteasome together with its biogenesis machinery. Crucially, the same module additionally contains canonical upstream regulators of the ubiquitin–proteasome system: UBC, the polyubiquitin precursor that supplies the ubiquitin pool for proteasomal substrate targeting^36^, and HSPA8, an Hsc70 chaperone that triages misfolded substrates toward proteasomal degradation^37,38^. The co-clustering of structural subunits with their assembly factors and functional regulators distinguishes this substruc-ture from a mere co-expression group: RICE groups these genes by participation in a shared regulatory program, not by similarity of expression alone. GAPDH also clusters here, consistent with its emerging non-glycolytic roles in proteostasis under oxidative stress^39^. A related proteostasis module recurs in RPE1 cells (Figure S5D), where the 20S subunit PSMB5 and the 19S ATPase PSMC5 co-cluster with protein-folding and stress-response factors: the mitochondrial co-chaperonin HSPE1, the oxidative-stress regulator PTMA^40^, and the tubulin folding cofactor TBCA^41^. The recurrence of proteasome subunits within a coherent proteostasis grouping across two independent cell lines indicates that the recovered structure reflects conserved regulatory organization rather than a cell-line-specific artifact.

To assess whether these clusters captured biologically meaningful signals, we performed Gene Ontology enrichment analysis^42,43^ across all clusters. Among significantly enriched terms (Fisher’s exact test with Benjamini–Hochberg correction, *p <* 0.01), we report the top 15 ranked by odds ratio in the “Biological Process” and “Molecular Function” categories (Figure 6E). Of the seven clusters enriched for at least one term, three showed enrichment in both categories. Enriched molecular function terms were unique to each cluster, and redundancy in biological process terms was minimal, with only three terms shared between two clusters. A similar pattern was observed in RPE1 cells (Figure S5E), where enriched terms were distinct across both categories.

The enriched biological processes are strongly centered on core essential functions, including mitotic spindle organization, chromosome segregation, and cell cycle checkpoint regulation, while molecular function terms highlight RNA polymerase activity, RNA binding, and ribosome-associated processes . These results are consistent with well-established categories of essential genes, reflecting both proliferative machinery and fundamental transcriptional and translational programs. Importantly, rather than arising from a generic list of essential genes, these enrichments emerge from the structure of the inferred network, suggesting that the proposed method captures biologically meaningful organization. The presence of coherent functional modules further indicates that the estimated DAG reflects structured dependencies among genes, providing evidence that the inferred relationships are not purely statistical but align with known biological systems.

### Revealing condition-specific immune regulatory signals in melanoma cells

We studied immune regulatory networks across conditions using a pooled Perturb-CITE-seq dataset of melanoma cells^44^. In this dataset, cells were co-cultured with autologous tumor-infiltrating lymphocytes (TILs), treated with interferon (IFN)-*γ*, or maintained under control conditions, to probe mechanisms of cancer immune evasion. The data consists of 182 genes after filtration.

RICE was run separately on each condition across 50 bootstrap replicates, and we selected edges with consistent sign (positive or negative) in at least 80% of replicates for downstream analysis. Resampling was stratified by perturbation: the number of perturbed cells for each gene, together with the number of control cells, was preserved in every bootstrap sample, holding the perturbation design fixed across replicates. This step improves model stability and is motivated by the sparsity of the data: roughly 40% of genes have mean expression below 0.5, which destabilizes their negative binomial regression coefficients. As a consequence, an edge may be selected owing to the low signal-to-noise ratio in these genes rather than a genuine and stable effect. This limitation of magnitude-based selection becomes pronounced under high sparsity; because the bootstrap procedure mitigates it, we recommend this approach for similarly sparse datasets.

The inferred networks comprised 113, 110, and 102 edges in the IFN-*γ*, co-culture, and con-trol environments, respectively, characterized by per-gene in- and out-degree (Figure 7). STAT1 was a dominant out-hub under IFN-*γ* and co-culture but not control (out-degree 25 and 15), consistent with its tumour-suppressive role: STAT1 mediates the anti-proliferative and pro-apoptotic effects of interferons and drives MHC class I-dependent recognition by cytotoxic T cells^45^, and its silencing suppresses MHC expression as immune evasion^46^. The receptor-proximal signalling genes IFNGR1, IFNGR2, and JAK1 were out-hubs only under immune stimulation, recovering the ligand-gated activation of the IFN-*γ* cascade—whose disruption (e.g. JAK1/2 loss) confers checkpoint-blockade resistance^47,48^. In contrast, the antigen-presentation genes B2M and HLA-B were hubs across all three environments, matching the constitutive, IFN-*γ*-amplified MHC class I machinery: B2M is the invariant light chain obligate for surface MHC class I, and its loss is a recurrent evasion mechanism in melanoma^49,50^. That immune stimulation concentrates IFN-*γ* pathway genes among the hubs while antigen-presentation effectors remain central throughout shows the inferred networks recover the canonical, context-specific JAK/STAT antigen-presentation program^44^.

**Figure 7:**
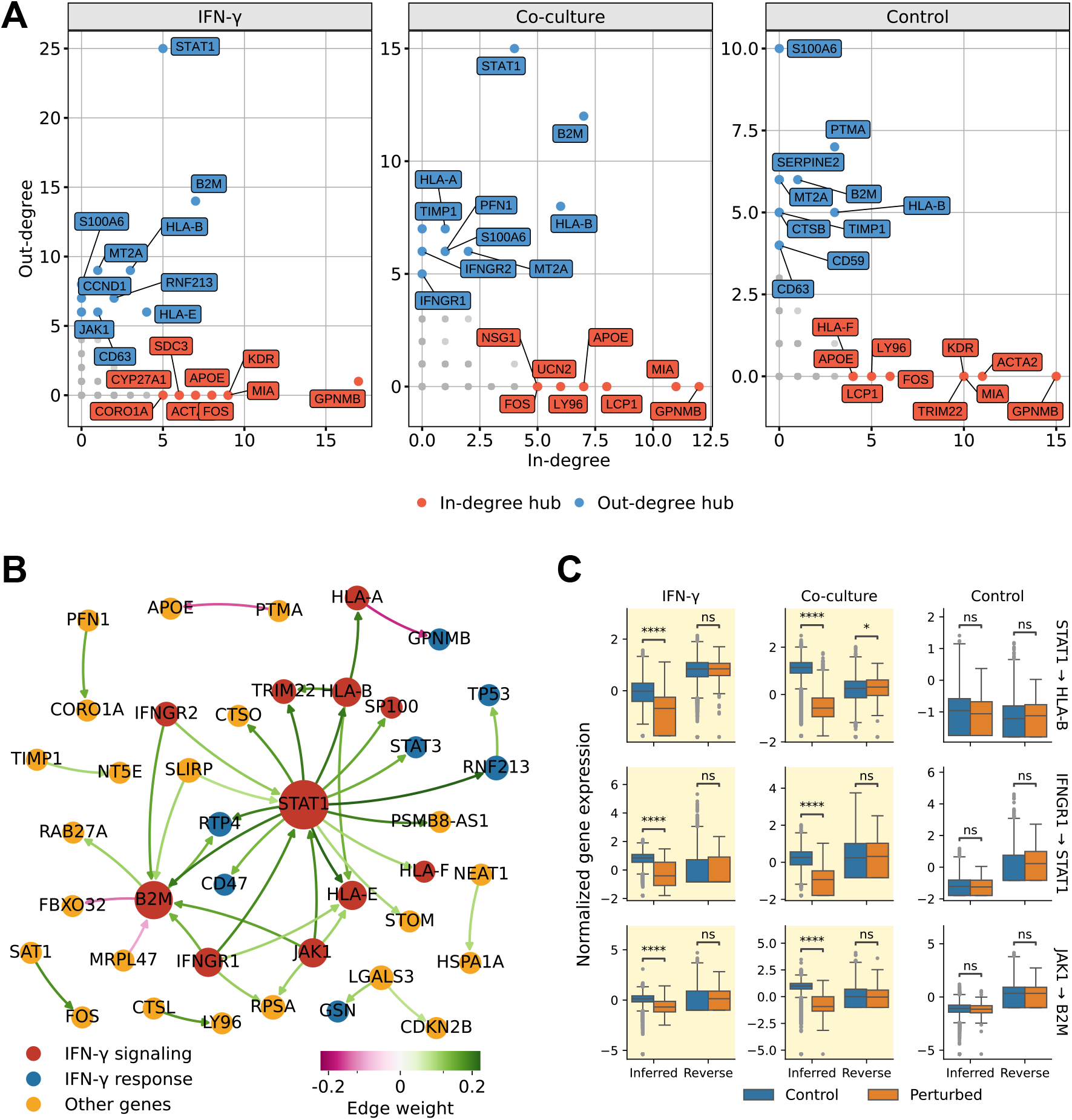
De novo discovery of JAK–STAT immune signaling in melanoma cells under IFN-*γ* or autologous TIL stimulation using RICE. Edges shown appear in at least 80% of 50 bootstrap replicates. **(A)** The top 10 genes by in-degree and out-degree in the inferred network across the three environments: IFN-*γ*, co-culture with TIL, and control. **(B)** Subgraph of edges present under IFN-*γ* and co-culture with TIL but absent from control. Edge color indicates the average edge weight across bootstrap samples in which the edge is selected. Node color indicates whether a gene is an IFN-*γ* signaling gene, an IFN-*γ*-regulated gene, or other (see Methods for definitions). Node size reflects degree within the subgraph. **(C)** Boxplots of normalized target-gene expression, comparing unperturbed versus source-gene-perturbed cells across all three environments, for three source–target pairs: STAT1 → HLA-B, IFNGR1 → STAT1, and JAK1 → B2M. Both the inferred and reverse directions are shown, and *p*-values are from the Mann–Whitney *U* test. A yellow background indicates that the edge was selected by RICE in the corresponding environment.

We next examined edges present under both IFN-*γ* and co-culture but absent from control (Figure 7B). A coherent core of the IFN-*γ* signalling cascade was shared across the stimulated environments: IFNGR1, IFNGR2, and JAK1 →STAT1 and B2M;STAT1→ B2M, HLA-B, HLA-E, and HLA-F; and HLA-B→ HLA-A all appeared under IFN-*γ* and co-culture but not control. These orientations match the canonical cascade—IFNGR and JAK1 act upstream of STAT1, which drives the MHC class I genes B2M and the HLA heavy chains^51,52^—and were identical across the two independent stimulated conditions, indicating that the method reconstructs a stable, correctly oriented structure rather than condition-specific noise. The HLA-B→HLA-A orientation was further supported interventionally: perturbing HLA-B significantly shifted the HLA-A distribution (*p <* 10^*−*4^ in all environments, Mann–Whitney *U* test) with no significant reciprocal effect.

Edge directions were further verified in Figure 7C: for representative edges (STAT1→ HLA-B, IFNGR1→ STAT1, JAK1→ B2M), the inferred effect was significant under IFN-*γ* and co-culture but not control, and significant in the inferred but not the reverse direction, confirming both the condition-specificity and the orientation of the recovered structure (additional comparisons in Figure S6). Beyond the canonical pathway, RICE identified STAT1 as the source of many downstream IFN-*γ* genes in both stimulated environments: for example, CD47, an IFN-*γ*-inducible immune-modulatory gene regulated via the JAK-STAT1-IRF1 axis^53,54^, and RNF213, an interferon-stimulated gene induced by IFN-*γ* downstream of STAT1-IRF1^55,56^. These edges represent a de novo recovery of interferon-driven immune structure with correct directionality.

Beyond recovering known biology, RICE yields novel candidate findings in this dataset. Figure 7A shows that GPNMB and MIA serve as in-degree hubs across all conditions, both with established immunological roles: GPNMB is an IFN-*γ*-associated immunosuppressive glycoprotein that negatively regulates tumor response to immune checkpoint inhibitors^57^, and MIA is a secreted melanoma marker linked to suppression of anti-tumor immunity^58^. Although their expression shows no significant shift when the sources are perturbed, both are expressed extremely sparsely (mean *<* 0.1), which limits statistical power and explains the large *p*-values. Importantly, the corresponding edges recur in at least 80% of bootstrap replicates under our negative binomial model, marking them as stable candidates despite the lack of significance. Considering their established immunosuppressive roles, this motivates their nomination as putative regulators. We similarly highlight S100A6, which emerges as an out-degree hub across all environments. RICE also identifies HLA-A, HLA-B→ HLA-F in every environment; these perturbations did not reach significance either, which we again attribute to the low expression of HLA-F (mean expression around 0.4). We recommend all of these for further investigation.

Taken together, these results show that our method recovers not merely individual gene-to-gene edges but a coherent regulatory structure consistent with both the data and prior biology, demonstrating its utility for reconstructing complex networks. Moreover, by surfacing stable reg-ulatory candidates that conventional significance tests overlook, RICE goes beyond confirming known biology to nominate novel hypotheses for further investigation based on the recovered causal structure.

## DISCUSSION

Across both synthetic and real-world datasets, RICE consistently outperforms existing methods, and its accuracy remains stable across data distributions and intervention types where current approaches, typically optimized for a single setting, degrade. Our ablation analysis shows that the reduced control function is the source of this robustness: removing it leaves true signals intact but admits a substantial number of false edges under strong confounding, and the resulting loss of precision is exactly the failure mode that corrupts network reconstruction in real perturbation data. This establishes the reduced control function not as an incidental design choice but as the component that makes confounding-robust causal discovery feasible at network scale, at the cost of a modest and readily affordable reduction in statistical power.

RICE also accommodates high-MOI designs directly, by modeling the combined effects of multiple perturbations rather than reducing each cell to a single primary perturbation through the heuristic simplifications common in prior methods. This treatment is increasingly relevant as high-MOI screens become a practical necessity for genome-scale studies. Supported by GPU-accelerated optimization, RICE scales to datasets with hundreds of genes and hundreds of thousands of cells, making it suitable for the size of modern single-cell perturbation experiments. Several limitations remain. The current framework assumes that perturbation effects combine additively and does not explicitly model higher-order genetic interactions. The acyclicity constraint, while essential for valid causal interpretation and tractable optimization, precludes the recovery of feedback loops, which are known to occur in many regulatory systems^59,60^. RICE therefore recovers the feedforward backbone of the regulatory network, and extending it to admit cyclic structure is an important direction for future work. In addition, as a point estimator, RICE selects edges by coefficient magnitude for efficient large-scale optimization; extending it to settings where edge uncertainty is natively supported is a further direction for methodological development.

In conclusion, RICE provides a robust and scalable framework for causal network reconstruction in increasingly complex single-cell perturbation experiments. By making confounding-robust causal discovery practical at scale, and by recovering networks that both recapitulate known regulatory biology and nominate new candidate regulators, RICE offers a foundation for moving from cataloguing perturbation effects toward a causal understanding of gene regulation.

## RESOURCE AVAILABILITY

### Lead contact

Requests for further information and resources should be directed to and will be fulfilled by the lead contact, Hongzhe Li.

### Materials availability

This study did not generate new materials.

### Data and code availability

- The K562 and RPE1 data are publicly available at https://doi.org/10.25452/figshare.plus.20029387.
- The melanoma data was downloaded via scPerturb^61^ at http://zenodo.org/records/7041849.
- The code for model training, simulation and data analysis is provided at https://github.com/HowardGech/RICE.
- Any additional information required to reanalyze the data reported in this paper is available from the lead contact upon request.

## ACKNOWLEDGMENTS

This work was funded by National Institutes of Health via grant R01GM129781 and U01HG013841. The authors thank all members of the lab for their support.

## METHODS

### Method details

#### Conditional distribution with observational and interventional data

Consider a Perturb-seq dataset involving *p* genes. For each cell, we observe (***y, z, D***), where ***y*** ∈ℕ^*p*^ is the vector of raw gene expression UMI counts, ***z*** ∈ℝ^*d*^ represents observed confounders (for example, library size and batch ID), and ***D*** {0, 1}^*p*^ is the perturbation indicator vector, where *D*_*j*_ = 1 indicates that gene *j* is perturbed and *D*_*j*_ = 0 otherwise. In addition, we allow for the presence of unmeasured confounders ***z***_um_.

Because linear relationships are rarely appropriate for count data, we assume that the influence between genes follows a univariate nonlinear monotone transformation *g*. Applying this transformation yields the variable ***x*** ∈ ℝ^*p*^, where *x*_*j*_ = *g*(*y*_*j*_). We further assume that the *p* genes form a DAG, such that each gene *j* is influenced only by its parent genes denoted by Pa(*j*). The conditional distribution of ***y*** is specified by

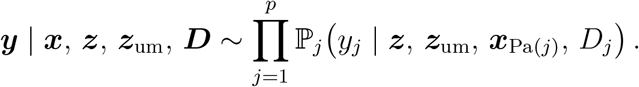

Under this framework, all non-perturbed genes retain an observational form, while perturbed genes adopt an intervention-modified conditional distribution.

Interventions in Perturb-seq can be incorporated at two levels of granularity. Soft interventions modify the conditional distribution of the perturbed gene by introducing a distribution shift, while still allowing it to be influenced by its parents. In contrast, hard interventions fully over-write upstream regulation: parent effects on the perturbed gene are removed, and the gene’s expression depends solely on confounders and the intervention effect. Both intervention types integrate seamlessly into our modeling and optimization framework; the underlying parameterization of the regulatory matrix and the DAG constraint remain identical, and only the regression structure—and thus the gradients—differ during optimization.

In this study, we model each conditional gene-expression distribution using a generalized linear model, where confounders, parent genes, and interventions contribute additively to the expression of gene *j*:

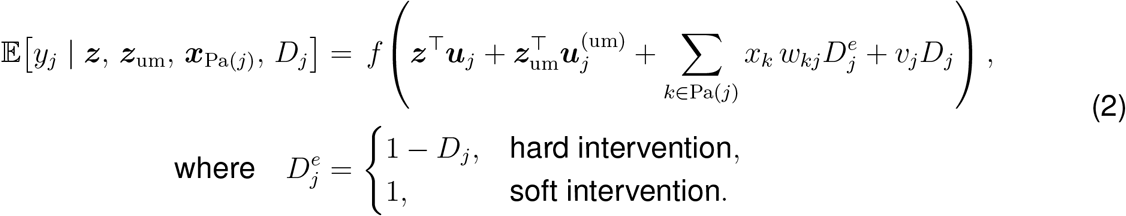

Here, ***u***_*j*_ and 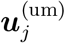 denote the coefficients for the confounders, *v*_*j*_ measures the intervention effect, and *D*_*j*_ is the perturbation indicator. The function *f* (·) is the link function determined by the assumed data distribution. The parameter *w*_*kj*_ captures the direct causal effect of gene *k* on gene *j*. The matrix ***W*** := (*w*_*kj*_)_1*≤k,j≤p*_ captures the causal relationships and is constrained to follow a DAG structure, ensuring a valid causal interpretation.

#### Reduced control function approach for unmeasured confounding

A common challenge in estimating causal gene networks is the presence of unmeasured confounding ***z***_um_. Because ***z***_um_ cannot be directly conditioned on, estimates of the causal effects ***W*** may be biased due to unblocked backdoor paths between genes. To address this issue, we leverage the exogenous nature of Perturb-seq interventions. Because sgRNAs (and thus the perturbation indicators ***D***) are assigned independently of unmeasured confounders, they serve as valid instrumental variables. Therefore, we adopt a reduced control function approach to proxy the influence of ***z***_um_.

The reduced control function approach proceeds in two stages. In the first stage, we isolate variation in the parent genes that is independent of unmeasured confounders. For each gene *k*, we model its transformed expression *x*_*k*_ as a function of the observed covariates ***z*** and the perturbation indicator *D*_*k*_. Let 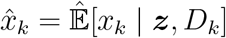 denote the predicted expression from this stage. We then define the first-stage residual as

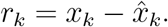

This residual *r*_*k*_ captures the component of variation in *x*_*k*_ that is not explained by observed confounders or the perturbation effect, and is therefore primarily driven by unmeasured confounders ***z***_um_ as well as intrinsic biological noise.

In the second stage, we augment the model for each gene by including the residuals of all other genes as additional covariates. The surrogate conditional expectation under the control function formulation replaces the original regression structure and is given by

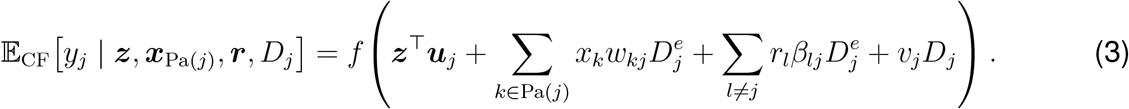

Here, *β*_*lj*_ denotes the coefficient of the control function term from gene *l* to gene *j*. By conditioning on the residuals of all other genes, we control for variations not associated with perturbations, including those from unmeasured confounding. This formulation isolates the directed causal influence among genes, ensuring that the DAG structural constraint and the causal interpretation of the weight matrix ***W*** remain valid even in the presence of unmeasured confounding.

Instead of the full control-function method, which computes 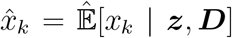, the reduced approach regresses *x*_*k*_ only on its own perturbation *D*_*k*_, excluding all other perturbations. To understand why this formulation remains robust in practice, we rewrite the terms inside the parenthesis of the right-hand side in (3) as

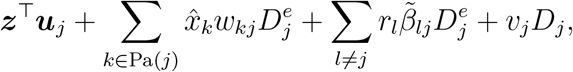

where 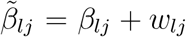. Because 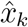 is orthogonal to *r*_*k*_, the coefficient *w*_*kj*_ primarily captures the effect of 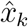 on *y*_*j*_. Under the reduced control-function approach, this effect is represented by *D*_*k*_ after conditioning on ***z*** and other genes, which is the standard basis for identifying whether gene *k* has a direct causal effect on gene *j* when *D*_*k*_ is treated as an instrumental variable. By contrast, replacing 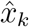with 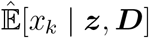makes the estimation of *w*_*kj*_ depend on the full perturbation vector ***D***. When the data violate model assumptions (such as the DAG or instrumental variable), the perturbation of other genes can induce false signals, therefore influencing the estimation of true causal effects. The identifiability results with the exponential link function are detailed in Note S2.

#### Constrained optimization and GPU acceleration

Under the control function formulation, the log-likelihood for many commonly used count distributions can be written as

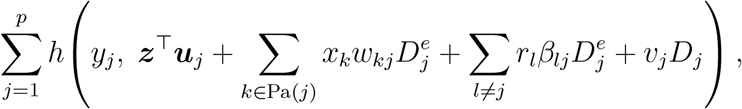

where *h*(·) is a smooth loss function.

Suppose we observe *n* cells, producing data matrices ***Y*** ∈ ℕ^*n×p*^, ***X*** ∈ℝ^*n×p*^, ***Z*** ∈ℝ^*n×d*^, and ***D*** ∈{0, 1}^*n×p*^. Let ***U*** = (***u***_1_, … , ***u***_*p*_), ***v*** = (*v*_1_, … , *v*_*p*_)^*⊤*^, ***B*** = (*β*_*lj*_)_1*≤l,j≤p*_, and ***R*** = (*r*_*ij*_); define ***D***^*e*^ as in (2). Maximizing the (regularized) log-likelihood is then equivalent to solving

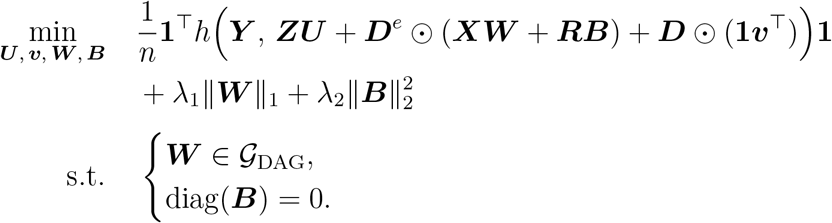

Here, ⊙denotes the Hadamard product, and *G* _DAG_ the set of matrices whose support corresponds to a DAG. The loss *h* is applied element-wise. An *l*_1_ penalty on ***W*** promotes sparsity in the inferred network, improving recovery in high-dimensional settings. An *l*_2_ penalty on ***B*** is introduced to mitigate the typically high correlation between ***X*** and ***R***, which would otherwise lead to unstable estimates.

Enforcing diag(***B***) = 0 is straightforward. However, directly imposing the DAG constraint is computationally prohibitive: the search space contains 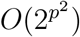 possible graphs, rendering exhaustive enumeration infeasible. Traditional algorithms require *O*(2^*p*^) operations in the worst case, which quickly becomes impractical as *p* grows. Bayesian approaches attempt to explore the DAG space using Markov Chain Monte Carlo (MCMC) moves that add, remove, or rewire edges, but these strategies also scale poorly for modern perturbation datasets involving thousands of genes.

To overcome these challenges, we adopt the framework of differentiable DAG learning, which converts the combinatorial problem of causal structure learning into continuous optimization through smooth acyclicity constraints. These formulations are smooth, differentiable, and provably equivalent to the exact DAG constraint, and several have been shown to recover both short- and long-range causal dependencies effectively. We therefore incorporate a smooth acyclicity constraint into our framework and solve the resulting constrained problem using an augmented Lagrangian approach. Table S1 provides a detailed comparison of three commonly used constraints.

A key advantage of this framework is that both the loss function and the acyclicity constraint are fully differentiable. This property enables the use of modern gradient-based optimizers, such as Adam, which are well suited to the nonconvex optimization landscape arising in this setting. The core computations of gradients—including matrix multiplications, element-wise operations, and those required by the acyclicity constraint (for example, matrix inversion, exponentiation or log-determinant evaluations)—are highly parallelizable and map naturally to modern GPU architectures. As a result, the method scales to large Perturb-seq datasets and high-dimensional gene networks, where GPU acceleration can provide substantial computational gains over traditional CPU-based causal discovery approaches.

#### Negative binomial distribution for overdispersion

Although the proposed framework applies to a broad class of generalized linear models for count data, we adopt the negative binomial distribution as the default likelihood because it provides a good empirical fit to single-cell RNA-seq measurements. In particular, gene expression counts from single-cell experiments typically exhibit substantial *overdispersion*, where the variance exceeds the mean, a phenomenon that cannot be adequately captured by the Poisson model. The NB distribution naturally accommodates overdispersion by introducing an additional dispersion parameter that allows the variance to vary independently of the mean.

We use the following parameterization of the NB distribution:

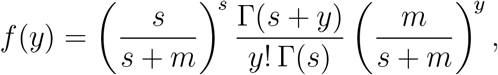

where Γ(·) denotes the Gamma function, *m >* 0 is the mean parameter, and *s >* 0 is the inverse dispersion parameter controlling the level of overdispersion. The corresponding negative log-likelihood for a single observation is

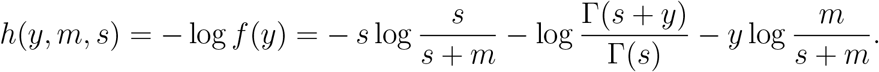

Under this parameterization, the distribution has mean *m* and variance *m* + *m*^2^*/s*, allowing the variance to exceed the mean when *s* is finite. An attractive property of this formulation is that as *s*→ ∞, the distribution converges to a Poisson distribution with mean *m*. The NB model can therefore be viewed as a flexible extension of the Poisson model that explicitly captures overdispersion while retaining closed-form likelihood and gradient expressions, enabling efficient optimization within our framework.

To incorporate the NB model into the causal network formulation, we parameterize both the mean and the dispersion using log links:

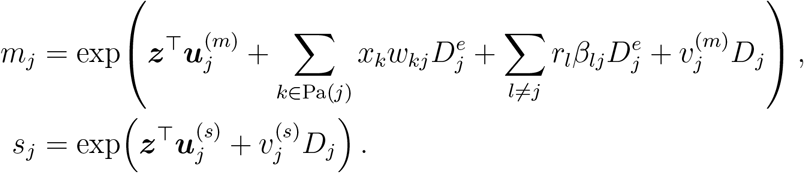

Here the mean parameter *m*_*j*_ is governed by the causal gene network while accounting for variation induced by unmeasured confounders through the control function terms. In contrast, the dispersion parameters *s*_*j*_ are assumed to be gene-specific and independent of other genes. Introducing this additional set of parameters allows the model to accommodate heterogeneous variability across genes while preserving the causal structure encoded in the network weights.

The resulting optimization objective then becomes

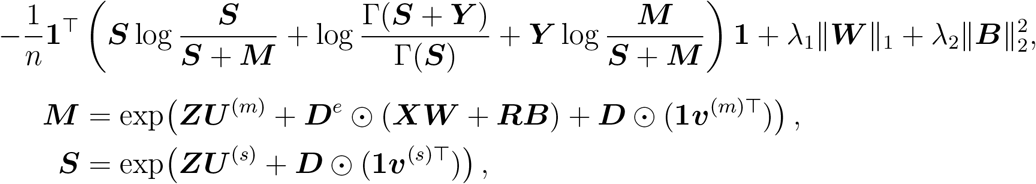

where the exponential is applied element-wise.

#### Augmented Lagrangian method and gradient computation

The proposed method is formulated within a general optimization framework using an augmented Lagrangian procedure:

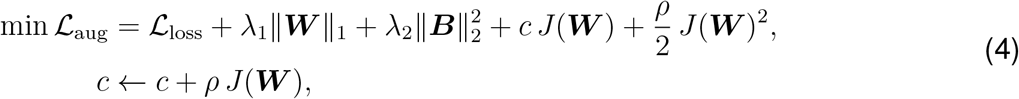

where *c* and *ρ* denote the dual variable and penalty coefficient, respectively, and *J*(***W***) represents the smooth DAG constraint. The loss function ℒ_loss_ depends on the model parameters, with gradients computed via the chain rule.

For the NB model, the loss admits a closed form, with gradients given by:

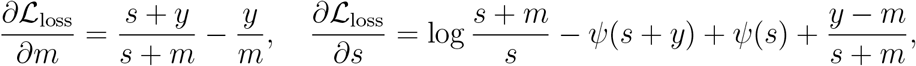

where *ψ*(·) is the digamma function.

#### Post-training sparsity recovery and hyperparameter selection

Although the *l*_1_ penalty in Eq. (4) encourages sparsity during optimization, the estimated weight matrix is typically not exactly sparse after training. In particular, for computational efficiency, we employ gradient-based optimizers such as Adam, which do not produce intrinsically sparse solutions. As a result, many entries remain nonzero but have very small magnitudes, forming a long tail of weak edge weights. In practice, these near-zero coefficients are difficult to distinguish from numerical shrinkage artifacts and can obscure the final causal graph if retained directly. We therefore apply a post-training thresholding step to recover a sparse network from the continuous edge-weight estimates.

Specifically, after model fitting, we compute the absolute values of all off-diagonal entries in 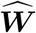,

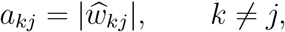

and sort them in decreasing order

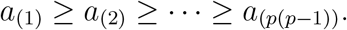

The resulting ordered sequence typically exhibits an initial regime corresponding to strong causal signals, followed by a flatter tail dominated by background noise. We identify the transition between these two regimes by detecting a change point in the sorted weight profile, for example using methods such as ruptures^62^ or kneed^63^, and take the corresponding magnitude as the threshold. Edges with absolute weight exceeding this threshold are retained, whereas the remaining edges are set to zero. An illustration of this procedure is shown in Figure S7A.

This procedure is fully data-driven and avoids the need to tune an edge-selection threshold using ground-truth graphs or validation labels, which are generally unavailable in real applications. Compared with imposing a fixed cutoff a priori, change-point-based thresholding adapts automatically to the scale of the learned weights in each fitted model and produces a sparse graph that is more readily interpretable from a biological perspective.

As shown in Figure S7B, this post hoc thresholding strategy performs comparably to oracle optimal thresholding across simulation replicates under both soft and hard interventions, suggesting that it provides an effective practical approximation when the true graph is unknown. We therefore use the adaptively thresholded graph as the final estimated causal network in all downstream analyses of real-world data.

The data-driven thresholding strategy further enhances model stability when the *l*_1_ penalty on the weight matrix (*λ*_1_) is small. While a low *λ*_1_ may theoretically select more false positives, these typically manifest as weak signals in the tail of the weight distribution. Consequently, the change-point detection algorithm tends to identify them as background noise and removes them automatically. In contrast, an excessively large *l*_1_ penalty can over-suppress true causal signals, making them indistinguishable from noise and leading to their unintended removal during thresholding. This is evident in the sensitivity analysis in Figure S7C, where higher *λ*_1_ values result in a clear decrease in TPR.

While we observed that increasing the *l*_2_ penalty for the control function (*λ*_2_) generally improves performance by reducing linear correlation, we caution against using an arbitrarily large *λ*_2_. As *λ*_2_→ ∞, the model effectively reverts to a version without the reduced control function and may therefore fail to account for unmeasured confounding. Given the inherent heterogeneity of single-cell screens—including differences in experimental platforms, cell types, and perturbation strengths—a single set of hyperparameters is unlikely to be optimal across all settings. We therefore recommend starting with a relatively small *λ*_1_ (e.g., 10^*−*4^) and a moderate *λ*_2_ (e.g., 0.01), and then refining these parameters based on the results of the initial fit.

#### Simulation details

The simulations in this work followed the model described in (2), while varying the data-generating distribution (*f*), the dimension of unmeasured confounding ***z***_um_, the structure and magnitude of the causal graph ***W*** , the perturbation indicator matrix ***D***, and the perturbation effect ***v***. The default setting consisted of 10,000 cells, 100 genes, and 400 true causal effects, with a negative binomial data-generating distribution and an Erdős–Rényi topology for the support of ***W*** . In the very-large-scale setting, we generated 100,000 cells, 500 genes, and 2,000 true causal effects. True causal effects were randomly sampled from [−1, −0.5] ∪ [0.5, 1] (or [−0.5, −0.3] ∪ [0.3, 0.5] under weak causality). Each element of the perturbation effect vector ***v*** was randomly sampled from [−3,−2] (or [−1.5,−1] under weak perturbation) to mimic CRISPRi-mediated inhibition. For the Poisson log-normal distribution, the standard deviation of the latent Gaussian component was fixed at 0.5.

All settings included an intercept term (*z* = 1) with coefficient −1. For the negative binomial distribution, the log-dispersion parameter log(*s*) also included an intercept term with coefficient 1; upon perturbation, an additional effect sampled from [−1, 1] (or [−0.5, 0.5] under weak perturbation) was added to the log-dispersion. Unmeasured confounders ***z***_um_ were drawn independently from a standard Gaussian distribution, with coefficients ***u***^(um)^ randomly sampled from [−1, 1]. To avoid unrealistically large or sparse counts, we adjusted the gene-wise log-mean expression vectors: we first shifted each vector by a constant such that its maximum value remained below 7.5 and then further shifted the vector to ensure its mean value remained above 0.5.

In the simulation with exclusion-restriction violations, we randomly select 5% of gene pairs to carry hidden effects. For the source gene *j* in each such pair, we define

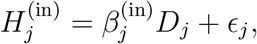

and for each target gene *k* we define S_*k*_ as the set of its source genes, and

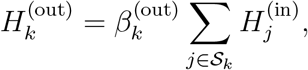

where the coefficients 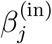and 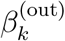 are each sampled uniformly from [−3, 3], and the noise terms *ϵ*_*j*_ are sampled uniformly from [0, 0.3]. For each *k*, the term 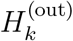 is then added to the argument of the link function in equation (2).

In low-MOI settings, 10% of samples were designated as control cells, and the remaining samples were partitioned into equal-sized groups and randomly assigned a single perturbation. In high-MOI settings, 10% of samples were again retained as controls, whereas the remaining samples were assigned a fixed number of random perturbations corresponding to the specified multiplicity of infection.

#### Training configuration

All numerical studies were conducted on an internal compute server equipped with Intel Xeon Gold 6542Y CPUs and NVIDIA H200 GPUs with 141 GB of VRAM. The Multi-Instance GPU

(MIG) feature was enabled on this platform, allowing the physical GPUs to be partitioned into compute instances ranging from a minimum of 16 GB to the maximum full-device capacity of 141 GB.

To ensure fair comparisons in model runtime benchmarking, CPU-based methods were strictly allocated 4 CPU cores, whereas GPU-based methods were assigned a single, unpartitioned physical GPU. For all other analyses, we maintained the 4-core CPU configuration but allocated the smallest MIG GPU instance that satisfied the memory requirements of each specific task to improve resource utilization.

RICE was implemented in Python with GPU-accelerated linear algebra via CuPy^64^, trained using the Adam optimizer with *β*_1_ = 0.9, *β*_2_ = 0.99, and a learning rate of 10^*−*4^. Unless otherwise specified, the regularization hyperparameters were set to *λ*_1_ = 10^*−*4^ and *λ*_2_ = 0.01. The augmented Lagrangian parameters were initialized at *ρ* = 1 and *c* = 0. For better convergence, *ρ* was updated using a multiplicative factor of 10 whenever the DAG constraint failed to decrease by at least 50% between successive iterations.

For baseline methods, the same hard-thresholding procedure was applied to the results of dotears. For inspre, the optimal model was selected via the authors’ recommended cross-validation, while for IBCD, we retained gene pairs that exhibited a posterior frequency greater than 0.95 across the MCMC samples, utilizing the default setting of 3,900 iterations.

#### Pre- and post-processing of K562 and RPE1 cells

The K562 and RPE1 results were derived from the CRISPRi Perturb-seq functional genomics maps generated by Replogle et al.^19^. Specifically, we used data from the essential-wide screens for model illustration and benchmarking. The raw K562 dataset initially contained 310,385 cells and 8,563 genes, with perturbations targeting 2,057 genes. To obtain a high-quality analytical dataset, we applied the following filtering criteria: (i) perturbations assigned to fewer than 100 cells were removed; (ii) genes with perturbation-effect *z*-scores below 5 were excluded; and (iii) genes with mean expression levels below 0.5 were filtered out. After preprocessing, the final K562 dataset contained 175,894 cells and 811 genes. The same filtering procedure was applied to the RPE1 cell line, which initially contained 247,914 cells, 8,749 genes, and 2,393 targeted perturbations. After filtering, the final RPE1 analytical dataset comprised 86,173 cells and 374 genes.

For both datasets, the library size was calculated as the total UMI count per cell, adjusted by the scale factor of the corresponding GEM group, and normalized by the total number of sequenced genes. This library size, along with the mitochondrial read percentage and one-hot-encoded GEM group identifiers, were included as observed covariates. To further account for latent technical factors and increase statistical power, we also included the top 50 principal components (PCs) derived from the set of excluded genes as additional covariates. To prevent these PCs from capturing downstream regulatory signals—which could lead to unwanted mediation effects or biased estimates—we regressed out the perturbation indicators from the PCs prior to their inclusion in the model. For other linear-model-based methods, we applied the Frisch– Waugh–Lovell theorem by first regressing out the covariates and then using the residuals as model inputs, thereby isolating causal effects not explained by observed confounders.

The input expression levels *x*_*ij*_ were computed as:

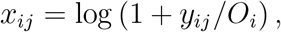

where *y*_*ij*_ represents the raw sequencing reads for gene *j* in cell *i*, and *O*_*i*_ denotes the total UMI count for that cell. Finally, we applied standard normalization to *x* to ensure that the inferred weights capture the relative magnitudes of the causal signals.

Following model optimization, we applied the kneed algorithm^63^ with a sensitivity parameter of *S* = 2.5 to determine the data-driven threshold for the weight matrix.

For external validation of inferred TF–target gene regulations, we integrated recorded interactions from all knowledge bases into a single unified regulatory network. An inferred gene pair (*j, k*) was considered annotated if a directed path existed from the upstream gene *j* to the downstream gene *k* within this consolidated network, and a reverse pair was considered annotated if a directed path existed from *k* to *j*. Pairs with directed paths in both directions were classified as bidirectional.

#### Pre- and post-processing of melanoma cells

The melanoma dataset was obtained from the multimodal pooled Perturb-CITE-seq screens of Frangieh et al.^44^. The raw data comprised 218,331 cells and 23,712 genes spanning 248 distinct perturbations. Although the reported data exhibited a high-MOI structure, this was not intended in the experimental design, and our exploratory analysis suggested that many detected guides likely reflected ambient RNA noise rather than functional gene knockouts. We therefore used the perturbation assignments provided by the original authors and treated the data as low-MOI to ensure model stability.

Given the relatively small number of perturbed genes, we adopted a more relaxed filtering strategy to retain sufficient data. We kept all genes with mean expression above 0.01 and at least 50 assigned cells per environment, yielding a final set of 182 genes. After filtering, the dataset comprised 75,213, 62,772, and 49,572 cells for the IFN-*γ*, co-culture, and control conditions, respectively. To account for technical and biological variation, the model included library size as well as mitochondrial and ribosomal read percentages as covariates. All other downstream procedures—PC calculation for excluded genes, data normalization, and weight-matrix thresholding—followed the same protocol as the analyses on the previous datasets.

We defined IFN-*γ* signaling genes as those in the Reactome “Interferon gamma signaling” gene set^65^. IFN-*γ* regulated genes were defined as those in the GO biological process “response to type II interferon” gene set^42^, the Hallmark “interferon gamma response” gene set^66^, or the Interferome database (type II interferon, melanoma cells)^67^.

#### Quantification and statistical analysis

Significance thresholds throughout the figures are indicated by asterisks: ^*\**^*p <* 0.05, ^*\*\**^*p <* 0.01, ^*\*\*\**^*p <* 0.001, and ^*\*\*\*\**^*p <* 10^*−*4^. Boxplots show the first quartile, median, and third quartile, with whiskers extending to the minimum and maximum values. Markers in strip plots and line plots denote the mean across all individual measurements. The specific statistical test, definition of error bars, and any further conventions are reported in the corresponding figure legend.

To evaluate causal discovery performance, we treat the presence of a true causal relationship between a pair of genes as a positive and the absence of such a relationship (or a reverse causal relationship) as a negative. Let TP, FP, TN, and FN denote the numbers of true positives, false positives, true negatives, and false negatives, respectively. Positive predictive value (PPV) and true positive rate (TPR) are then defined as

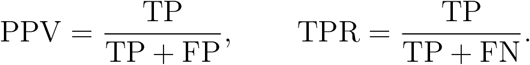

The Matthews correlation coefficient (MCC) is computed as in standard binary classification:

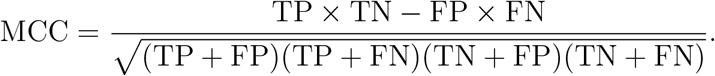

The structural Hamming distance (SHD) is defined as the minimum number of edge additions, deletions, or reversals required to transform the inferred causal graph into the ground-truth network.

Total and direct causal effects in the Perturb-seq data were estimated by linear regression with heteroskedasticity-robust (HC3) standard errors. Principal components derived from excluded genes (used as covariates during model fitting) were not included in these post hoc analyses. The effect of perturbing gene *k* on the expression of gene *j* was estimated as

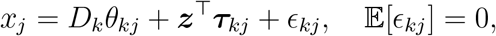

where *θ*_*kj*_ is the perturbation effect, ***τ***_*kj*_ is the vector of coefficients for the observed confounders ***z***, and *ϵ*_*kj*_ is the error term.

To isolate the direct causal effect, we additionally conditioned on the estimated parent set of gene 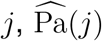, excluding gene *k* when it was itself a parent. The direct effect was then defined through

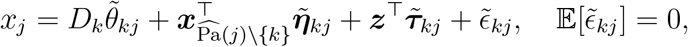

where 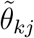 is the direct causal effect, 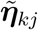captures the contributions of the remaining parent genes, and 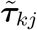 accounts for the observed confounders. Statistical significance of the total and direct effects was assessed using the *p*-values of *θ*_*kj*_ and 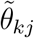, respectively.

We use linear regression here for several reasons. First, total causal effects in linear SEMs are conventionally estimated by linear regression, so this follows standard practice for the quantity being computed. Second, it provides a fair benchmark against the competing methods, which themselves assume linear structures; estimating effects with a GLM while comparing to linear-SEM methods would confound the comparison. Third, the choice is empirically immaterial: our preliminary results show that the GLM estimates differ little from the linear-model estimates. Finally, linear regression is substantially more efficient to compute than a GLM at this scale. We retain the GLM in equation (2) where it matters—modeling raw counts within RICE’s estimation framework—while using linear regression for the standalone effect estimates, where its assumptions are appropriate and its results are equivalent in practice.

To quantify the enrichment of significant regulatory interactions within the inferred network, we computed the odds ratio as

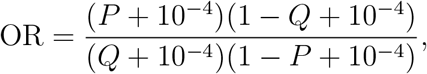

where *P* is the proportion of inferred edges that are statistically significant, and *Q* is the proportion of statistically significant edges among a randomly sampled set of edges of the same size. The constant 10^*−*4^ is added for numerical stability.

