## Supplementary Material for "Robust causal gene network estimation for large-scale single-cell perturbation screens using reduced control function"

Table S1: **Comparison of smooth acyclicity constraints for DAG learning.**

| | **Trace-exponential** | **Log-determinant** | **Trace-binomial** |
|---|---|---|---|
| **Constraint** | $\mathrm{tr}(\exp(W \odot W)) - p = 0$ | $\log \lvert sI - W \odot W \rvert - p \log s = 0$ | $\sum_{j=1}^{p} \mathrm{tr}((W \odot W)^j)/j = 0$ |
| **Gradient** | $2W \odot \exp(W^\top \odot W^\top)$ | $2W \odot (sI - W^\top \odot W^\top)^{-1}$ | $2W \odot \sum_{j=0}^{p-1} (W^\top \odot W^\top)^j$ |
| **Complexity** | $O(p^3)$ | $O(p^3)$ | $O(p^3 \log p)$ |
| **Advantage** | Easy to optimize | Avoids long-range cycles | Avoids long-range cycles |
| **Limitation** | Limited in representing long-range cycles | Requires additional assumptions | Computationally expensive |

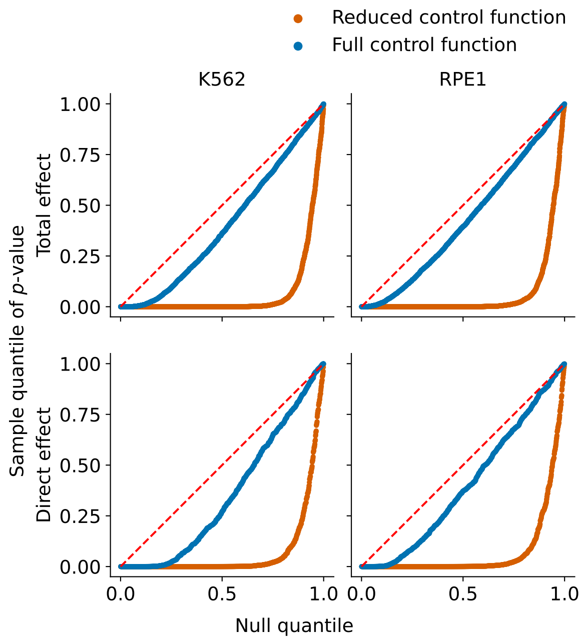


Figure S1: **Quantile-quantile plots of the $p$-values for total and direct causal effects in K562 and RPE1 cells, with gene networks estimated using the reduced and full control function framework.** The full control function approach yields much worse result than the reduced control function in this setting.

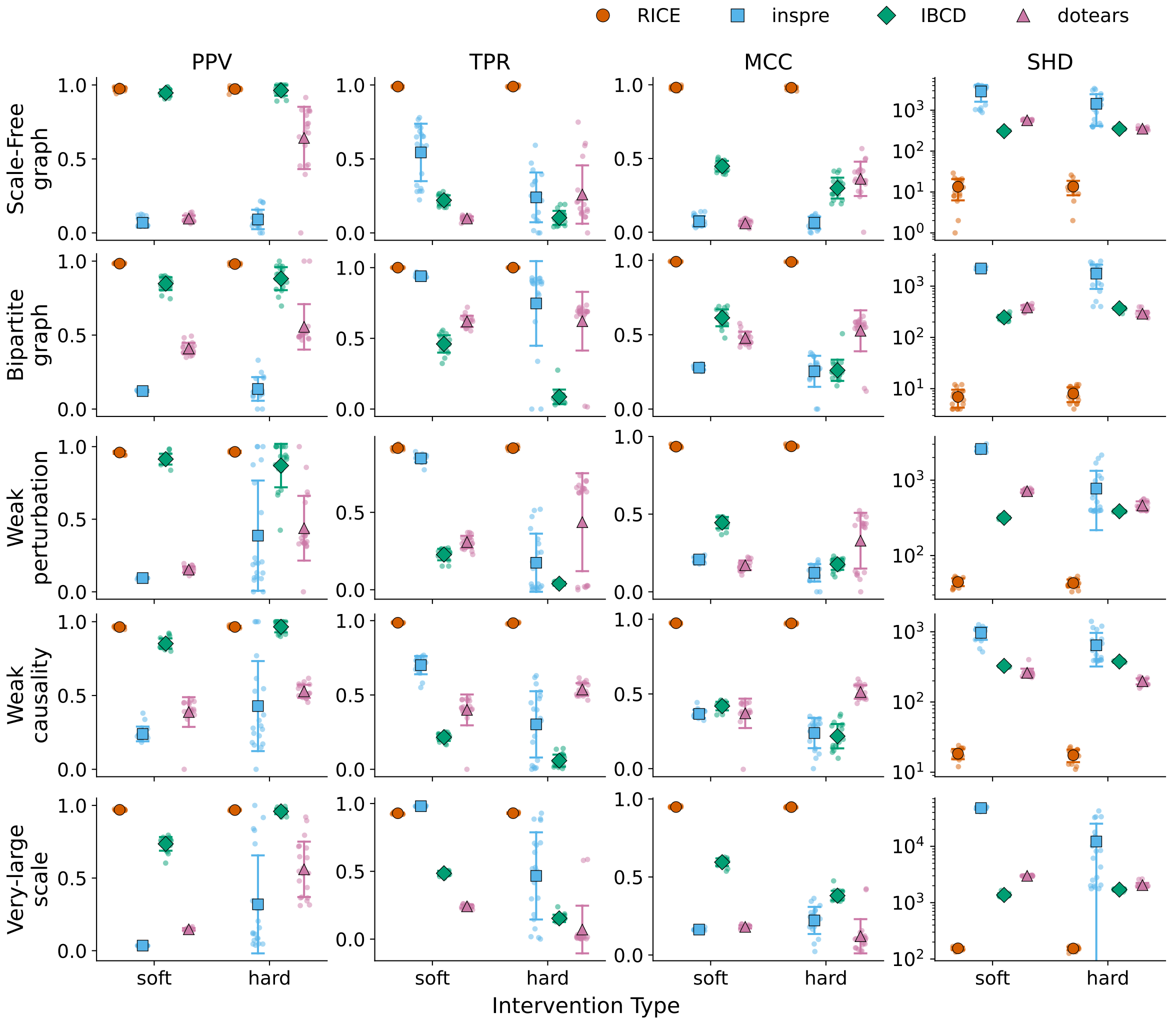


Figure S2: **Additional simulation results under scale-free graphs, bipartite graphs, weak perturbation, weak causality and very-large graph scale.** The data generation process was described in the simulation details section in the main manuscript.

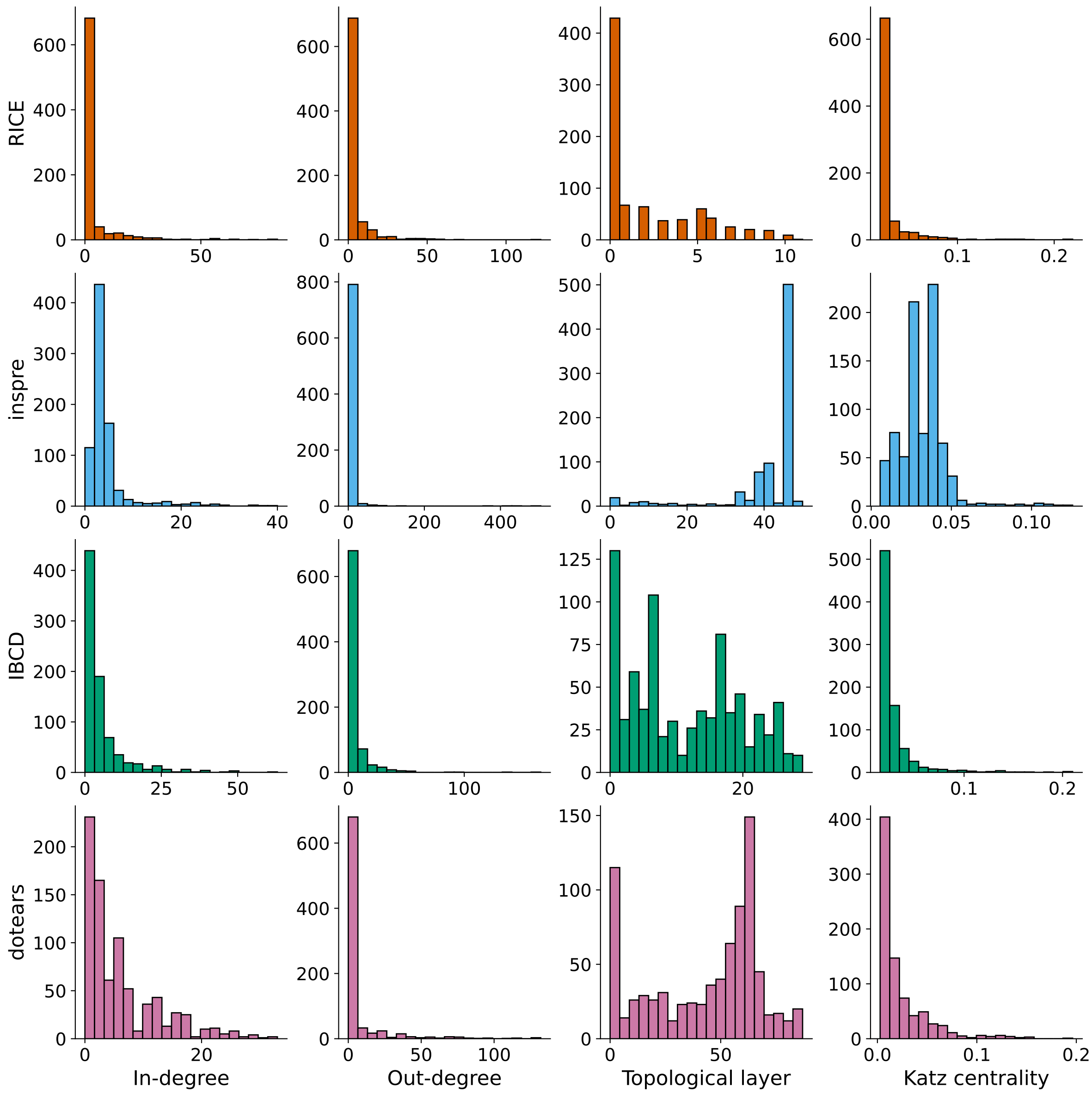


Figure S3: **Properties of causal gene networks in K562 cells estimated by four methods**. Katz centrality was computed with hyperparameters $\alpha = 0.1$ and $\beta = 1$.

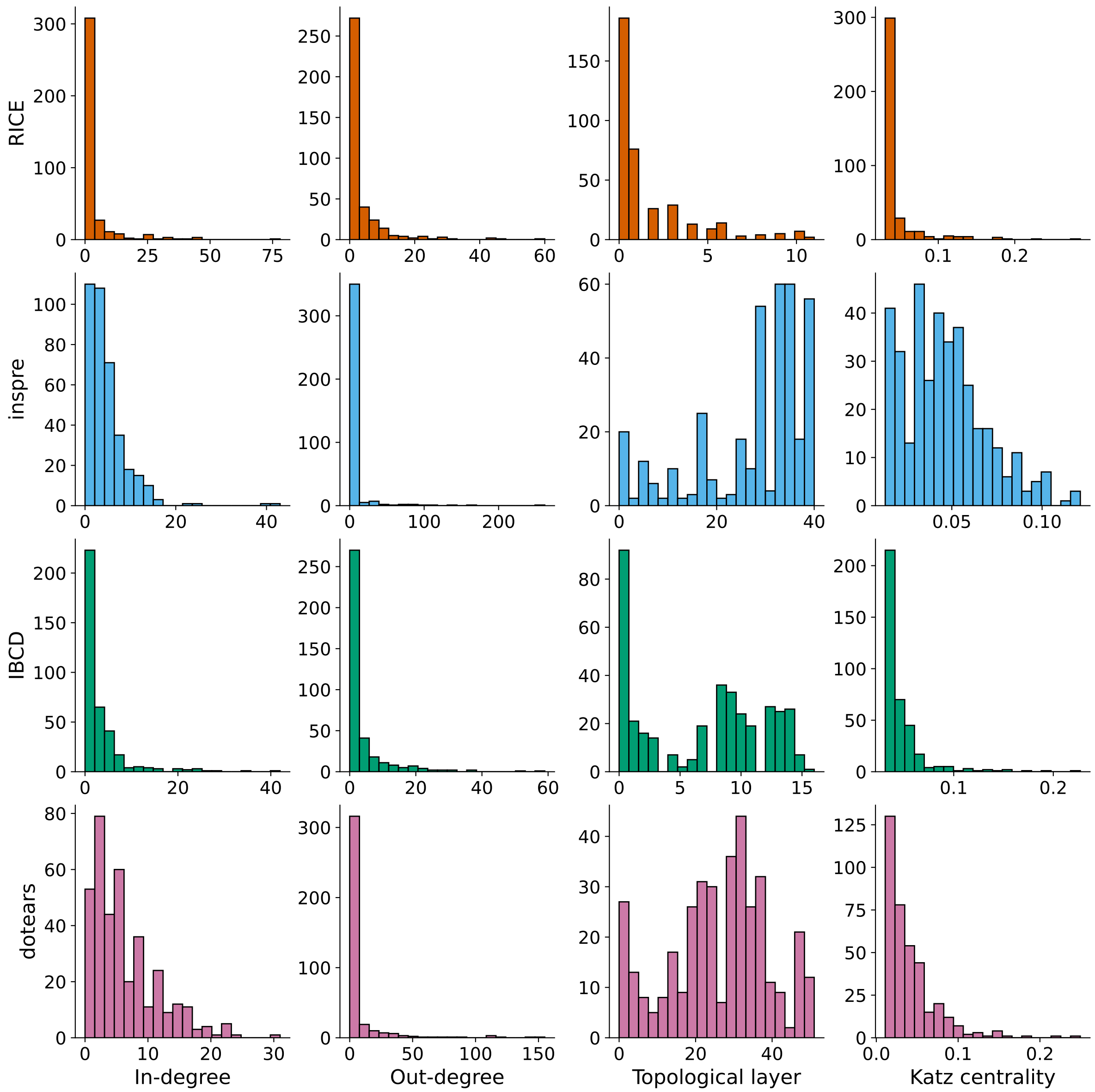


Figure S4: **Properties of causal gene networks in RPE1 cells estimated by four methods.** Katz centrality was computed with hyperparameters $\alpha = 0.1$ and $\beta = 1$.

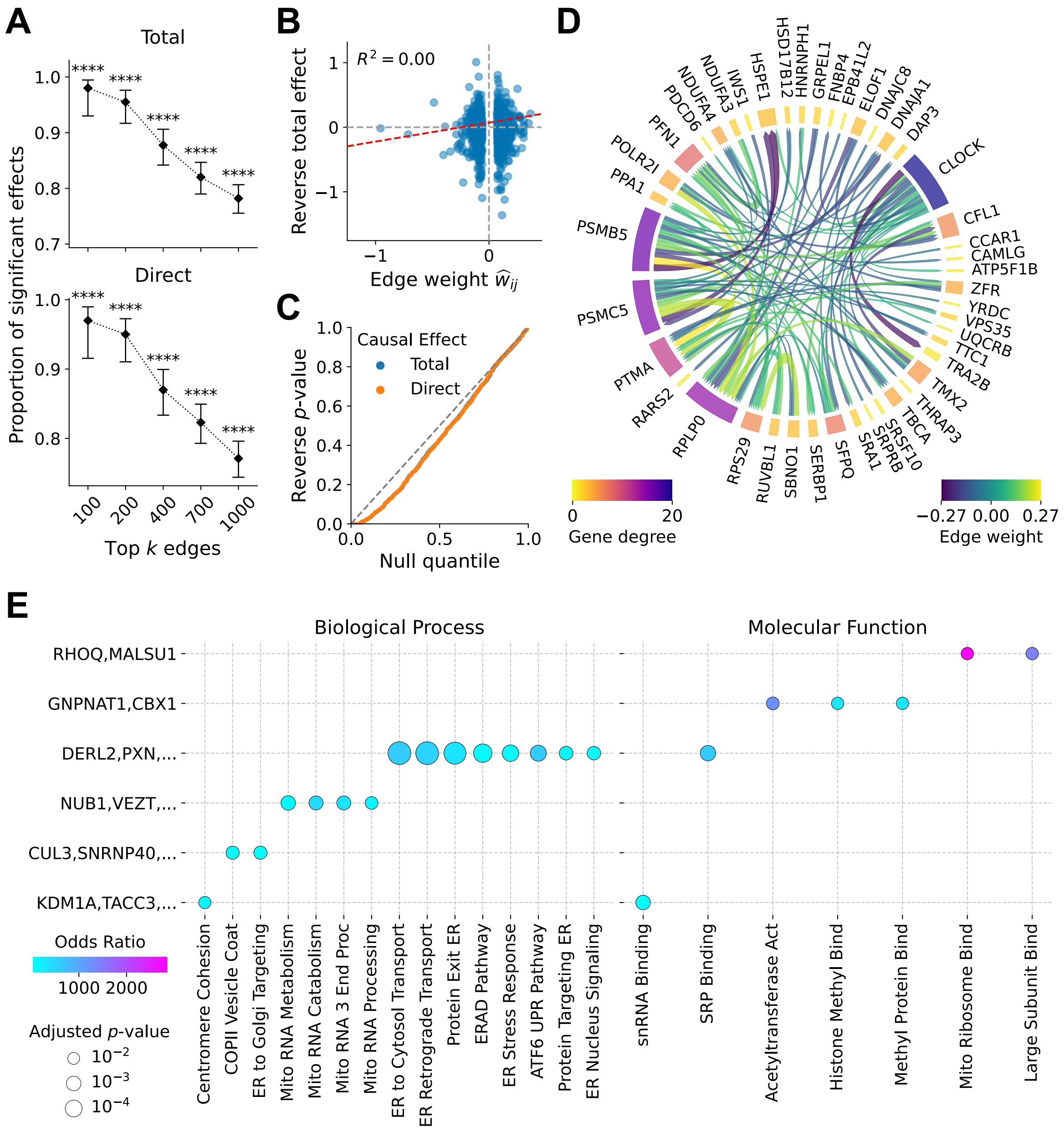


Figure S5: **Analysis of RICE-identified causal gene network in RPE1 cells.** **(A)** Proportions of top-ranked edges (from top 1,000 to top 100) exhibiting significant total or direct causal effects ($t$-test, $p < 0.01$). The proportion of validated effects increases with higher edge importance. Error bars denote 95% Wilson binomial confidence intervals. **(B)** Correlation between total causal effects of reversed regulator–target pairs and inferred edge weights. Red dashed lines represent linear regression fits; $R^2$ denotes the coefficient of determination. **(C)** Quantile–quantile plots of $p$-values for reversed regulator–target pairs compared against the theoretical uniform distribution under the null. **(D)** Representative gene cluster with 44 genes identified via Louvain community detection. **(E)** Gene Ontology enrichment analysis across identified clusters for "Biological Process" and "Molecular Function" categories. The top 15 terms ranked by odds ratio among those with Benjamini–Hochberg adjusted $p < 0.01$ are shown. Only clusters containing at least one enriched term are displayed.

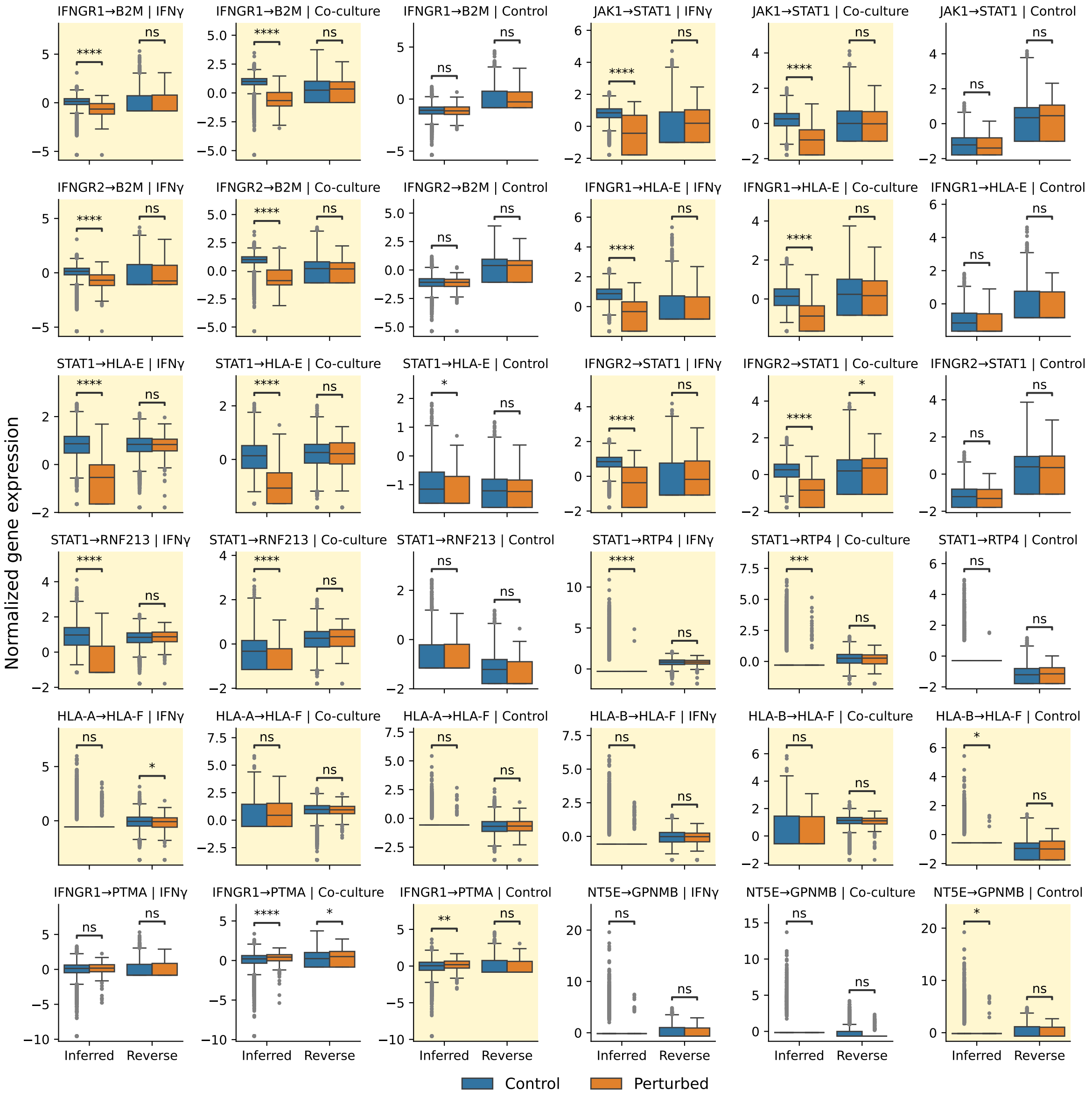


Figure S6: **Boxplots of normalized target-gene expression, comparing unperturbed versus source-gene-perturbed cells across all three environments, for additional source–target pairs.** Both the inferred and reverse directions are shown, and $p$-values are from the Mann–Whitney $U$ test. A yellow background indicates that the edge was selected by RICE in the corresponding environment.

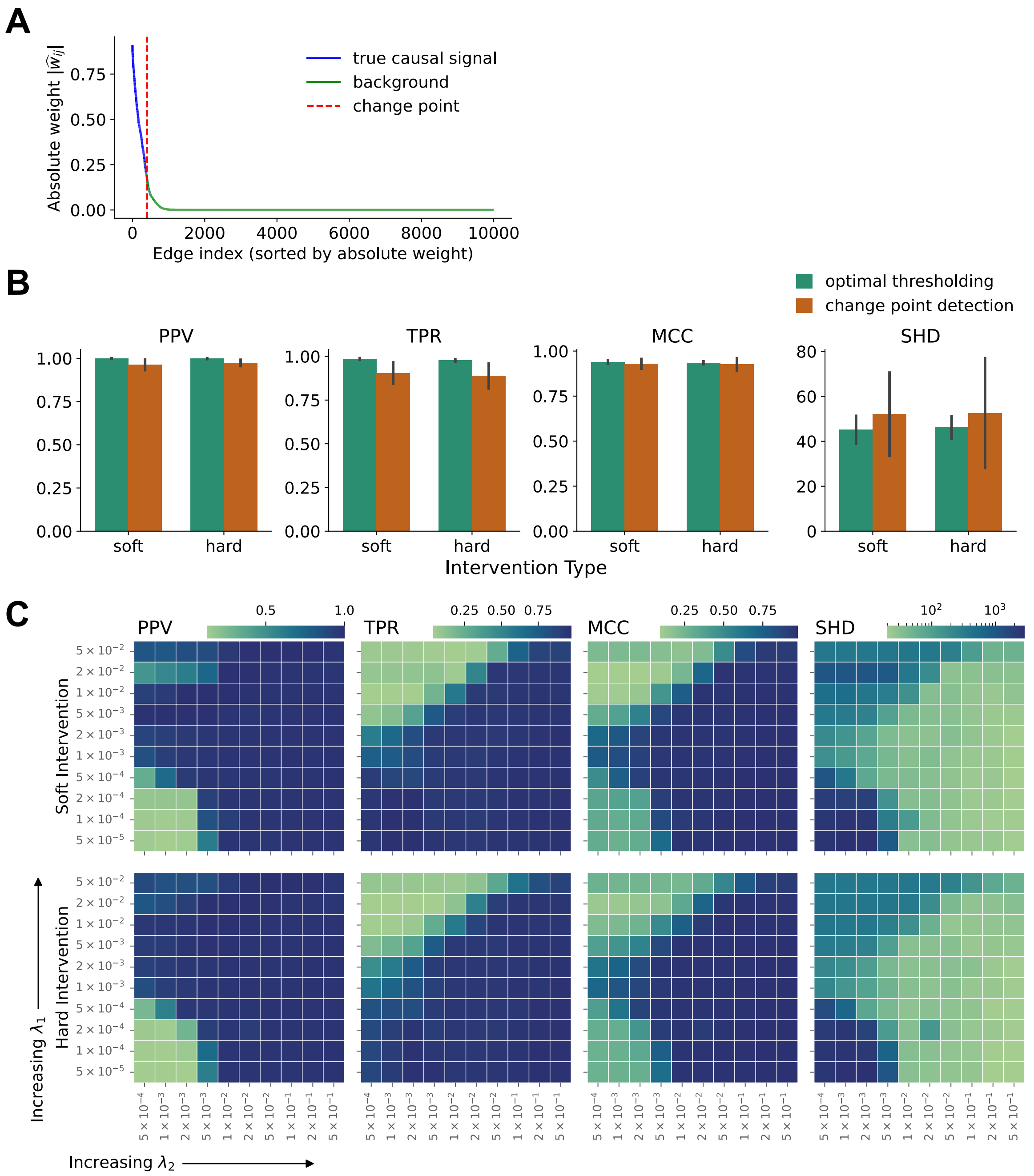


Figure S7: **Change-point-based thresholding and hyperparameter sensitivity analysis.** **(A)** Representative example of estimated network weights after thresholding by change-point detection. RICE produces a long tail of near-zero edge weights, which is removed by thresholding. The detected change point defines a threshold that effectively separates true causal signals from background noise. **(B)** Comparison of optimal thresholding and change-point-based thresholding across 20 replicates. Optimal thresholding denotes the best result over the threshold grid $\{0.1, 0.2, 0.3, 0.4, 0.5\}$. Error bars indicate the standard deviation.
**(C)** Sensitivity analysis of RICE hyperparameters. $\lambda_1$ ranged from $5 \times 10^{-5}$ to 0.05, while $\lambda_2$ ranged from $5 \times 10^{-4}$ to 0.5.

### Note S1 More probabilistic models

We consider another two probabilistic models: the Poisson distribution and the Poisson Log-Normal (PLN) distribution. Each model induces a different link function, which in turn determines the corresponding loss function $\mathcal{L}_{\text{loss}}$ in the optimization objective.

**Poisson distribution** We begin with the Poisson model, whose loss takes the form

$$\mathcal{L}_{\text{loss}}(\lambda) = \lambda - k \log \lambda. \tag{1}$$

The gradients are simple:

$$\frac{\partial \mathcal{L}_{\text{loss}}}{\partial \lambda} = 1 - \frac{k}{\lambda}. \tag{2}$$

A key strength of the Poisson DAG model is that it is *identifiable* from observational data alone(Park and Raskutti, 2015), meaning the full DAG structure can be recovered without interventions. In contrast, most graphical models can recover only a *Markov equivalence class* from observational data, leaving edge directions undetermined(He et al., 2015). This makes the Poisson model appealing when perturbation data are scarce or unavailable. However, this identifiability relies on the strict Poisson property that the mean equals the variance. In single-cell RNA-seq data, where overdispersion is prevalent, this assumption is violated, limiting the practical applicability of a pure Poisson model.

**Poisson Log-Normal distribution** To address overdispersion, we next consider the PLN model, in which the Poisson rate parameter follows a log-normal distribution:

$$\mathcal{L}_{\text{loss}}(\lambda) = -\log \int_{-\infty}^{\infty} \exp(kx - e^x)\, d\Phi(x \mid \log \lambda, \sigma^2), \tag{3}$$

where $\Phi(\cdot \mid \log \lambda, \sigma^2)$ is the cumulative distribution function of a normal distribution with mean $\log \lambda$ and variance $\sigma^2$. Unlike the Poisson model, the PLN distribution naturally accommodates overdispersion, with the extent of variance inflation controlled by $\sigma^2$. This flexibility has motivated its widespread adoption in recent single-cell gene expression analysis (Chiquet et al., 2019; Ge and Li, 2025). However, the loss function has no closed-form expression, and its gradient is likewise intractable, requiring numerical approximation during optimization.

Because the PLN distribution is defined as a mixture, a natural approximation strategy is to use the expectation–maximization (EM) framework. Introducing a latent variable $\boldsymbol{l} \in \mathbb{R}^p$, the PLN likelihood admits the hierarchical representation

$$\begin{gathered} \boldsymbol{y} \mid \boldsymbol{x}, \boldsymbol{l}, \boldsymbol{z}, \boldsymbol{D} \sim \prod_{j=1}^{p} \text{Poisson}\big(y_j \mid \lambda_j e^{l_j}\big)\,, \\ \boldsymbol{l} \mid \sigma^2 \sim \mathcal{N}(\boldsymbol{0},\, \sigma^2 \boldsymbol{I}), \end{gathered} \tag{4}$$

where $\mathcal{N}(\boldsymbol{\mu}, \boldsymbol{\Sigma})$ denotes a multivariate normal distribution with mean vector $\boldsymbol{\mu}$ and covariance matrix $\boldsymbol{\Sigma}$. Here, $\lambda_j$ corresponds to the exponential of all additive effects, including covariates, perturbation and effects from upstream genes. The resulting expected negative log-likelihood, up to an additive constant, is given by

$$\mathbb{E}_{\boldsymbol{l} \sim P(\cdot \mid \hat{\boldsymbol{\lambda}})} \sum_{j=1}^{p} \left( -\frac{1}{2} \log \sigma^2 + \frac{l_j^2}{2\sigma^2} + \lambda_j e^{l_j} - y_j (l_j + \log \lambda_j) \right), \tag{5}$$

where $\hat{\boldsymbol{\lambda}}$ denotes the current parameter estimates. Since the posterior distribution $P(\boldsymbol{l} \mid \widehat{\boldsymbol{U}}, \hat{\boldsymbol{v}}, \widehat{\boldsymbol{W}}, \widehat{\boldsymbol{B}})$ remains intractable, we replace it with a mean-field variational approximation:

$$\begin{gathered} \widetilde{P}(\boldsymbol{l} \mid \hat{\boldsymbol{\lambda}}) = \underset{Q \in \mathcal{Q}_{MF}}{\arg\min}\, D_{\text{KL}}\Big(Q \,\|\, P(\boldsymbol{l} \mid \hat{\boldsymbol{\lambda}})\Big)\,, \\ \mathcal{Q}_{MF} = \left\{ \mathcal{N}(\boldsymbol{\mu},\, \text{diag}(e^{\boldsymbol{\tau}})) : \boldsymbol{\mu} \in \mathbb{R}^p,\ \boldsymbol{\tau} \in \mathbb{R}^p \right\}, \end{gathered} \tag{6}$$

where $D_{\mathrm{KL}}$ denotes the Kullback–Leibler (KL) divergence and $\mathrm{diag}(e^{\boldsymbol{\tau}})$ is a diagonal covariance matrix containing the variational variances. Under this approximation, the surrogate loss function becomes

$$\tilde{\mathcal{L}}_{\mathrm{loss}}(\lambda) = \mathbb{E}_{l\sim\widetilde{P}}\left(\lambda e^{l} - k(l + \log\lambda)\right) = \lambda e^{\mu+\tau/2} - k\log\lambda. \tag{7}$$

Maximizing the variational log-likelihood in (5) also yields an update for the hyperparameter $\sigma^2$. Setting the gradient with respect to $\sigma^2$ to zero leads to the update

$$\sigma^2 \leftarrow \frac{1}{p}\left(\|\boldsymbol{\mu}\|_2^2 + \|e^{\boldsymbol{\tau}}\|_2^2\right), \tag{8}$$

which enables the variance parameter to be inferred directly from the data rather than specified beforehand. By providing an initial value for $\sigma^2$, the algorithm can iteratively refine it in a fully data-driven manner.

The surrogate loss function for the PLN model also has analytical gradients:

$$\frac{\partial\tilde{\mathcal{L}}_{\mathrm{loss}}}{\partial\lambda} = \exp(\mu + e^{\tau}/2) - \frac{k}{\lambda}. \tag{9}$$

The next step is to derive the variational parameters $\mu$ and $\tau$. Expanding the KL divergence yields

$$D_{KL} = \sum_{j=1}^{p}\left(\frac{1}{2}\log\frac{\sigma^2}{e^{\tau_j}} - \frac{1}{2} + \frac{e^{\tau_j} + \mu_j^2}{2\sigma^2} + \hat{\lambda}_j\exp(\mu_j + e^{\tau_j}/2) - y_j\mu_j\right), \tag{10}$$

which is separable across the $p$ genes. Each coordinate can therefore be optimized independently through $p$ parallel univariate subproblems. Letting $\mathcal{L}_{KL}$ denote the univariate objective, its gradients are

$$\frac{\partial\mathcal{L}_{KL}}{\partial\mu} = \frac{\mu}{\sigma^2} + \hat{\lambda}\exp(\mu + e^{\tau}/2) - y, \quad \frac{\partial\mathcal{L}_{KL}}{\partial\mu} = -\frac{1}{2} + \frac{e^{\tau}}{2\sigma^2} + \frac{\hat{\lambda}}{2}\exp(\mu + e^{\tau}/2 + \tau), \tag{11}$$

and the Hessian takes the form

$$\nabla^2_{\mu,\tau}\mathcal{L}_{KL} = \begin{pmatrix} \frac{1}{\sigma^2} + \hat{\lambda}\exp(\mu + e^{\tau}/2) & \frac{\hat{\lambda}}{2}\exp(\mu + e^{\tau}/2 + \tau) \\ \frac{\hat{\lambda}}{2}\exp(\mu + e^{\tau}/2 + \tau) & \frac{e^{\tau}}{2\sigma^2} + \frac{\hat{\lambda}}{2}\exp(\mu + e^{\tau}/2 + \tau)\left(\frac{e^{\tau}}{2} + 1\right) \end{pmatrix}. \tag{12}$$

Each univariate subproblem is convex, so second-order Newton updates provide fast and stable convergence.

**Comparison of different models** A comparison across all three distributions is reported in Table 1, where $\mathrm{RICE}_{PS}$, $\mathrm{RICE}_{PLN}$, and $\mathrm{RICE}_{NB}$ denote RICE with Poisson, Poisson log-normal, and negative binomial likelihoods, respectively. We also include a Gaussian version of RICE, denoted $\mathrm{RICE}_{GS}$, as a baseline, and compare all variants against existing methods. Overall, methods based on count-valued models perform well, whereas methods that do not account for the count nature of the simulated data show substantially weaker performance.

Notably, $\mathrm{RICE}_{PS}$ and $\mathrm{RICE}_{PLN}$ show a moderate drop in performance when applied to negative binomial data, with precision, $F_1$ score, and MCC often falling below 90%. By contrast, $\mathrm{RICE}_{NB}$ consistently maintains performance close to 95% across nearly all metrics and data-generating distributions. This pattern is largely attributable to the underlying likelihood assumptions. The Poisson model is sensitive to overdispersion, while the Poisson log-normal model can accommodate overdispersion only when its magnitude is relatively homogeneous across samples. In contrast, the negative binomial model explicitly captures dispersion through a tunable parameter, making it flexible enough to encompass both alternatives as special cases.

Note S1 Table 1: **Mean of causal graph identification metrics on synthetic data.**

| Data Dist. | Method | Soft intervention | | | | Hard intervention | | | |
|---|---|---|---|---|---|---|---|---|---|
| | | PPV | TPR | $F_1$ | MCC | PPV | TPR | $F_1$ | MCC |
| Poisson | $\text{RICE}_{PS}$ | **99.5** | **99.5** | **99.5** | **99.5** | 99.4 | **99.5** | 99.5 | 99.5 |
| | $\text{RICE}_{PLN}$ | 99.3 | 99.4 | 99.3 | 99.3 | **99.6** | 99.5 | **99.5** | **99.5** |
| | $\text{RICE}_{NB}$ | 92.3 | 96.1 | 94.1 | 94.1 | 94.5 | 97.1 | 95.8 | 95.8 |
| | $\text{RICE}_{GS}$ | 31.8 | 39.8 | 35.3 | 34.8 | 36.6 | 45.2 | 40.4 | 40.0 |
| | inspre | 28.1 | 15.3 | 12.6 | 15.2 | 33.2 | 9.8 | 8.2 | 11.2 |
| | IBCD | 24.3 | 22.6 | 21.6 | 21.7 | 15.0 | 8.0 | 9.5 | 9.6 |
| | dotears | 34.2 | 44.0 | 37.4 | 37.4 | 46.8 | 56.1 | 51.0 | 50.7 |
| Poisson Log-Normal | $\text{RICE}_{PS}$ | 96.6 | 98.3 | 97.5 | 97.5 | 97.8 | 98.8 | 98.3 | 98.3 |
| | $\text{RICE}_{PLN}$ | **96.8** | **98.5** | **97.6** | **97.6** | **98.1** | **98.9** | **98.5** | **98.5** |
| | $\text{RICE}_{NB}$ | 94.4 | 97.6 | 95.9 | 95.9 | 97.2 | 98.6 | 97.9 | 97.9 |
| | $\text{RICE}_{GS}$ | 32.1 | 39.5 | 35.3 | 34.8 | 36.9 | 44.8 | 40.4 | 40.0 |
| | inspre | 37.6 | 13.8 | 11.0 | 15.0 | 36.7 | 9.3 | 6.2 | 10.2 |
| | IBCD | 22.6 | 19.5 | 19.5 | 19.4 | 11.3 | 7.1 | 8.0 | 7.7 |
| | dotears | 36.4 | 42.5 | 36.5 | 36.9 | 51.2 | 59.4 | 54.9 | 54.6 |
| Negative Binomial | $\text{RICE}_{PS}$ | 76.1 | 92.2 | 83.3 | 83.5 | 77.0 | 93.1 | 84.2 | 84.5 |
| | $\text{RICE}_{PLN}$ | 86.7 | 92.3 | 89.4 | 89.3 | 88.4 | 93.3 | 90.8 | 90.7 |
| | $\text{RICE}_{NB}$ | **98.2** | **98.7** | **98.5** | **98.4** | **98.2** | **98.6** | **98.4** | **98.4** |
| | $\text{RICE}_{GS}$ | 39.9 | 40.8 | 40.3 | 39.7 | 48.2 | 49.1 | 48.7 | 48.1 |
| | inspre | 16.6 | 6.7 | 3.9 | 5.4 | 15.0 | 5.7 | 2.4 | 3.9 |
| | IBCD | 14.1 | 12.9 | 12.5 | 12.2 | 6.9 | 4.2 | 4.9 | 4.4 |
| | dotears | 42.5 | 47.0 | 44.6 | 44.1 | 55.4 | 58.3 | 56.8 | 56.4 |

This simulation setting uses 5,000 samples, 100 genes, and 100 true causal links. Each result is averaged over 50 replicates and reported as a percentage (%). The best results are marked in bold.

### Note S2 Identifiability results: structural model and reduced control function

In this section, we separate the identifiability analysis into two parts.

The first part concerns the *structural model*. It establishes identifiability of the causal coefficient matrix under a relaxed first-stage assumption in which the expression of each gene may depend on the entire perturbation pattern $\boldsymbol{D}$. This result clarifies the population target of the model and does not rely on control-function adjustment.

The second part concerns the *reduced control-function formulation*. It shows that the same causal coefficient matrix remains identifiable when one works through the reduced residuals

$$r_i := x_i - \bar{m}_i(\boldsymbol{z}, D_i), \qquad \bar{m}_i(\boldsymbol{z}, D_i) := \mathbb{E}[x_i \mid \boldsymbol{z}, D_i].$$

Throughout, $\boldsymbol{x} = (x_1, \ldots, x_p)^\top$, $\boldsymbol{y} = (y_1, \ldots, y_p)^\top$, $\boldsymbol{D} = (D_1, \ldots, D_p)^\top \in \{0,1\}^p$, $\boldsymbol{z} \in \mathbb{R}^d$, and $\boldsymbol{z}_{\text{um}} \in \mathbb{R}^q$. The structural mean model is

$$\mathbb{E}[y_j \mid \boldsymbol{x}, \boldsymbol{z}, \boldsymbol{z}_{\text{um}}, D_j] = \exp\left(\boldsymbol{z}^\top \boldsymbol{\alpha}_j + \phi_j(\boldsymbol{z}, \boldsymbol{z}_{\text{um}}) + \sum_{k=1}^{p} x_k w_{kj}^0 D_j^e + v_j D_j\right), \tag{13}$$

where

$$D_j^e = \begin{cases} 1, & \text{soft intervention,} \\ 1 - D_j, & \text{hard intervention.} \end{cases}$$

#### Part I. Identifiability of the structural model

We first state assumptions that concern only the data-generating mechanism.

**A1 (Acyclicity).** $\boldsymbol{W}_0 = (w_{kj}^0)$ is the weighted adjacency matrix of a directed acyclic graph.

*Interpretation.* This defines the causal target as a DAG and excludes directed feedback loops.

**A2 (Exogeneity of perturbations).**

$$\boldsymbol{D} \perp \boldsymbol{z}_{\text{um}} \mid \boldsymbol{z}.$$

*Interpretation.* Conditional on observed covariates, the perturbation pattern is independent of the unmeasured confounders.

**A3 (Relaxed first-stage model).** For each gene $i$,

$$x_i = m_i(\boldsymbol{z}, \boldsymbol{D}) + h_i(\boldsymbol{z}, \boldsymbol{z}_{\text{um}}) + \varepsilon_i, \qquad m_i(\boldsymbol{z}, \boldsymbol{D}) = \mathbb{E}[x_i \mid \boldsymbol{z}, \boldsymbol{D}]. \tag{14}$$

*Interpretation.* The perturbation-driven component of $x_i$ is allowed to depend on the full perturbation pattern $\boldsymbol{D}$, not only on $D_i$. This allows, for example, perturbations of upstream regulators to affect downstream genes.

**A4 (Residual noise invariance).**

$$\mathbb{E}[\varepsilon_i \mid \boldsymbol{z}, \boldsymbol{z}_{\text{um}}, \boldsymbol{D}] = 0, \qquad \boldsymbol{\varepsilon} \mid (\boldsymbol{z}, \boldsymbol{z}_{\text{um}}, \boldsymbol{D}) \stackrel{d}{=} \boldsymbol{\varepsilon} \mid (\boldsymbol{z}, \boldsymbol{z}_{\text{um}}),$$

where $\boldsymbol{\varepsilon} = (\varepsilon_1, \ldots, \varepsilon_p)^\top$.

*Interpretation.* Conditional on $(\boldsymbol{z}, \boldsymbol{z}_{\text{um}})$, the residual noise is mean-zero and its distribution is not directly shifted by the perturbation design.

**A5 (First-stage identifiability).** The function

$$\boldsymbol{m}(\boldsymbol{z}, \boldsymbol{D}) := (m_1(\boldsymbol{z}, \boldsymbol{D}), \ldots, m_p(\boldsymbol{z}, \boldsymbol{D}))^\top$$

is identified from the joint distribution of $(\boldsymbol{x}, \boldsymbol{z}, \boldsymbol{D})$.

*Interpretation.* Since $m_i(\boldsymbol{z}, \boldsymbol{D}) = \mathbb{E}[x_i \mid \boldsymbol{z}, \boldsymbol{D}]$, this is a standard population identifiability condition for the first-stage conditional mean.

**Reduced representation of the structural model**

**Lemma 1.** *Under A2–A4,*

$$\mathbb{E}[y_j \mid \boldsymbol{z}, \boldsymbol{D}] = \exp\left(\sum_{k=1}^{p} m_k(\boldsymbol{z}, \boldsymbol{D})\, w_{kj}^0\, D_j^e\right) \Lambda_j(\boldsymbol{z}, D_j) \tag{15}$$

*for some measurable function $\Lambda_j$.*

*Interpretation.* After integrating out latent confounding and residual noise, the full perturbation pattern $\boldsymbol{D}$ enters the conditional mean only through the first-stage mean vector $\boldsymbol{m}(\boldsymbol{z}, \boldsymbol{D})$, together with a nuisance factor depending on whether the target gene $j$ itself is perturbed.

*Proof.* Substituting (14) into (13) gives

$$\begin{aligned}\mathbb{E}[y_j \mid \boldsymbol{x}, \boldsymbol{z}, \boldsymbol{z}_{\mathrm{um}}, D_j] = \exp\Bigg( & \boldsymbol{z}^\top \boldsymbol{\alpha}_j + \phi_j(\boldsymbol{z}, \boldsymbol{z}_{\mathrm{um}}) \\ & + \sum_{k=1}^{p} m_k(\boldsymbol{z}, \boldsymbol{D}) w_{kj}^0 D_j^e + \sum_{k=1}^{p} h_k(\boldsymbol{z}, \boldsymbol{z}_{\mathrm{um}}) w_{kj}^0 D_j^e \\ & + \sum_{k=1}^{p} \varepsilon_k w_{kj}^0 D_j^e + v_j D_j \Bigg).\end{aligned} \tag{16}$$

Taking conditional expectation given $(\boldsymbol{z}, \boldsymbol{z}_{\mathrm{um}}, \boldsymbol{D})$ yields

$$\begin{aligned}\mathbb{E}[y_j \mid \boldsymbol{z}, \boldsymbol{z}_{\mathrm{um}}, \boldsymbol{D}] = \exp\Bigg( & \boldsymbol{z}^\top \boldsymbol{\alpha}_j + \phi_j(\boldsymbol{z}, \boldsymbol{z}_{\mathrm{um}}) + \sum_{k=1}^{p} m_k(\boldsymbol{z}, \boldsymbol{D}) w_{kj}^0 D_j^e \\ & + \sum_{k=1}^{p} h_k(\boldsymbol{z}, \boldsymbol{z}_{\mathrm{um}}) w_{kj}^0 D_j^e + v_j D_j \Bigg) M_j(\boldsymbol{z}, \boldsymbol{z}_{\mathrm{um}}, D_j),\end{aligned} \tag{17}$$

where

$$M_j(\boldsymbol{z}, \boldsymbol{z}_{\mathrm{um}}, D_j) := \mathbb{E}\left[ \exp\left( \sum_{k=1}^{p} \varepsilon_k w_{kj}^0 D_j^e \right) \middle| \boldsymbol{z}, \boldsymbol{z}_{\mathrm{um}}, \boldsymbol{D} \right].$$

By A4, $M_j$ depends on $\boldsymbol{D}$ only through $D_j$. Integrating over $\boldsymbol{z}_{\mathrm{um}}$ conditional on $(\boldsymbol{z}, \boldsymbol{D})$ and using A2 yields (15). □

**Model identifiability under low-MOI**

**ML1 (Low-MOI support).** Conditional on $\boldsymbol{z}$, the support of $\boldsymbol{D}$ is contained in

$$\{\mathbf{0}, \boldsymbol{e}_1, \ldots, \boldsymbol{e}_p\},$$

where $\mathbf{0}$ is the all-zero vector and $\boldsymbol{e}_i$ is the $i$-th standard basis vector. Moreover,

$$\mathbb{P}(\boldsymbol{D} = \mathbf{0} \mid \boldsymbol{z}) > 0, \qquad \mathbb{P}(\boldsymbol{D} = \boldsymbol{e}_i \mid \boldsymbol{z}) > 0$$

on a set of positive probability for each $i$.

**ML2 (Invertibility of the low-MOI response matrix).** For each target node $j$, define

$$\boldsymbol{\Gamma}_j(\boldsymbol{z}) := \Big( m_k(\boldsymbol{z}, \boldsymbol{e}_i) - m_k(\boldsymbol{z}, \mathbf{0}) \Big)_{i \neq j,\ k \neq j}. \tag{18}$$

Assume $\boldsymbol{\Gamma}_j(\boldsymbol{z})$ is invertible on a set of positive probability.

**Theorem 1** (Model identifiability under low-MOI)**.** *Under A1–A5 and ML1–ML2, the matrix $\boldsymbol{W}_0$ is identifiable.*

*Intuition.* Each contrast between a single-perturbation state and the control state yields one linear equation in the unknown coefficients of the $j$-th column. The matrix $\boldsymbol{\Gamma}_j(\boldsymbol{z})$ collects these equations. Its invertibility guarantees uniqueness.

*Proof.* Fix $j$. For each $i \neq j$, both $\boldsymbol{D} = \boldsymbol{0}$ and $\boldsymbol{D} = \boldsymbol{e}_i$ satisfy $D_j = 0$, hence $D^e_j = 1$. By (15),

$$\mathbb{E}[y_j \mid \boldsymbol{z}, \boldsymbol{D} = \boldsymbol{0}] = \exp\left(\sum_{k=1}^{p} m_k(\boldsymbol{z}, \boldsymbol{0}) w^0_{kj}\right) \Lambda_j(\boldsymbol{z}, 0),$$

and

$$\mathbb{E}[y_j \mid \boldsymbol{z}, \boldsymbol{D} = \boldsymbol{e}_i] = \exp\left(\sum_{k=1}^{p} m_k(\boldsymbol{z}, \boldsymbol{e}_i) w^0_{kj}\right) \Lambda_j(\boldsymbol{z}, 0).$$

Taking the log-ratio gives

$$g_{ij}(\boldsymbol{z}) := \log \frac{\mathbb{E}[y_j \mid \boldsymbol{z}, \boldsymbol{D} = \boldsymbol{e}_i]}{\mathbb{E}[y_j \mid \boldsymbol{z}, \boldsymbol{D} = \boldsymbol{0}]} = \sum_{k=1}^{p} \big(m_k(\boldsymbol{z}, \boldsymbol{e}_i) - m_k(\boldsymbol{z}, \boldsymbol{0})\big) w^0_{kj}.$$

Since $w^0_{jj} = 0$,

$$g_{ij}(\boldsymbol{z}) = \sum_{k \neq j} \big(m_k(\boldsymbol{z}, \boldsymbol{e}_i) - m_k(\boldsymbol{z}, \boldsymbol{0})\big) w^0_{kj}.$$

Writing this system in vector form yields

$$\boldsymbol{g}_j(\boldsymbol{z}) = \boldsymbol{\Gamma}_j(\boldsymbol{z}) \boldsymbol{w}^{(-j)}_{\cdot j},$$

where

$$\boldsymbol{g}_j(\boldsymbol{z}) := (g_{ij}(\boldsymbol{z}))_{i \neq j}, \qquad \boldsymbol{w}^{(-j)}_{\cdot j} := (w^0_{kj})_{k \neq j}.$$

By A5, $\boldsymbol{\Gamma}_j(\boldsymbol{z})$ is identified from $(\boldsymbol{x}, \boldsymbol{z}, \boldsymbol{D})$, and $\boldsymbol{g}_j(\boldsymbol{z})$ is identified from $(y_j, \boldsymbol{z}, \boldsymbol{D})$. By ML2, $\boldsymbol{\Gamma}_j(\boldsymbol{z})$ is invertible on a set of positive probability, hence

$$\boldsymbol{w}^{(-j)}_{\cdot j} = \boldsymbol{\Gamma}_j(\boldsymbol{z})^{-1} \boldsymbol{g}_j(\boldsymbol{z})$$

is uniquely determined. Since $j$ was arbitrary, $\boldsymbol{W}_0$ is identifiable. □

**Model identifiability under high-MOI / multi-perturbation designs**

**MH1 (Availability of a target-unperturbed stratum).** For each $j$, there exists $d_j \in \{0, 1\}$ such that $D^e_j(d_j) = 1$ and

$$\mathbb{P}(D_j = d_j \mid \boldsymbol{z}) > 0$$

on a set of positive probability.

**MH2$_{ij}$ (Coefficient-wise non-collinearity of the first-stage mean).** Fix $j$ and $i \neq j$. Assume that there exists $d_j$ as in MH1 such that

$$\mathrm{Var}\Big(m_i(\boldsymbol{z}, \boldsymbol{D}) - \mathbb{E}[m_i(\boldsymbol{z}, \boldsymbol{D}) \mid \boldsymbol{z}, D_j = d_j, \{m_k(\boldsymbol{z}, \boldsymbol{D}), k \neq i, j\}] \;\Big|\; \boldsymbol{z}, D_j = d_j\Big) > 0 \tag{19}$$

on a set of positive probability.

**Theorem 2** (Model identifiability under high-MOI)**.** *Fix $j$. Under A1–A5, MH1, and MH2$_{ij}$ for all $i \neq j$, the $j$-th column of $\boldsymbol{W}_0$ is identifiable. If these conditions hold for every $j$, then $\boldsymbol{W}_0$ is identifiable.*

*Intuition.* Within a stratum where the target gene $j$ remains effectively unperturbed, the log conditional mean is an affine function of the first-stage mean vector. The non-collinearity condition guarantees that each coordinate of that vector contributes uniquely.

*Proof.* Fix $j$. Suppose another coefficient vector $\tilde{\boldsymbol{w}}_{\cdot j}$ yields the same model. By (15),

$$\sum_{k=1}^{p} m_k(\boldsymbol{z}, \boldsymbol{D})(w^0_{kj} - \tilde{w}_{kj})D^e_j + c_j(\boldsymbol{z}, D_j) = 0$$

almost surely for some measurable $c_j$. Restrict to a stratum $\{D_j = d_j\}$ with $D^e_j(d_j) = 1$. Since $w^0_{jj} = \tilde{w}_{jj} = 0$,

$$\sum_{k \neq j} m_k(\boldsymbol{z}, \boldsymbol{D})(w^0_{kj} - \tilde{w}_{kj}) + b_j(\boldsymbol{z}, d_j) = 0$$

almost surely given $(\boldsymbol{z}, D_j = d_j)$.

Fix $i \neq j$. Define

$$a_k := w^0_{kj} - \tilde{w}_{kj}, \qquad k \neq j.$$

Then

$$a_i m_i(\boldsymbol{z}, \boldsymbol{D}) + \sum_{k \neq i,j} a_k m_k(\boldsymbol{z}, \boldsymbol{D}) + b_j(\boldsymbol{z}, d_j) = 0$$

almost surely given $(\boldsymbol{z}, D_j = d_j)$. Let

$$\eta_{ij} := m_i(\boldsymbol{z}, \boldsymbol{D}) - \mathbb{E}[m_i(\boldsymbol{z}, \boldsymbol{D}) \mid \boldsymbol{z}, D_j = d_j, \{m_k(\boldsymbol{z}, \boldsymbol{D}), k \neq i, j\}].$$

Multiply the previous identity by $\eta_{ij}$ and take conditional expectation given $(\boldsymbol{z}, D_j = d_j)$. The terms involving $\{m_k(\boldsymbol{z}, \boldsymbol{D}), k \neq i, j\}$ and $b_j(\boldsymbol{z}, d_j)$ vanish, leaving

$$a_i\, \mathbb{E}[\eta^2_{ij} \mid \boldsymbol{z}, D_j = d_j] = 0.$$

By MH2$_{ij}$,

$$\mathbb{E}[\eta^2_{ij} \mid \boldsymbol{z}, D_j = d_j] > 0$$

on a set of positive probability, hence $a_i = 0$. Therefore

$$w^0_{ij} = \tilde{w}_{ij}.$$

Since $i \neq j$ was arbitrary, the $j$-th column is identifiable. Repeating this argument for all $j$ proves the final claim. □

#### Part II. Identifiability through the reduced control function

We now turn to the reduced control-function formulation. Here the objective is not merely to identify the structural model, but to show that the same causal coefficient matrix can be recovered when one works through the reduced residuals

$$r_i := x_i - \bar{m}_i(\boldsymbol{z}, D_i), \qquad \bar{m}_i(\boldsymbol{z}, D_i) := \mathbb{E}[x_i \mid \boldsymbol{z}, D_i].$$

The difficulty is that under the relaxed first-stage model A3,

$$x_i = m_i(\boldsymbol{z}, \boldsymbol{D}) + h_i(\boldsymbol{z}, \boldsymbol{z}_{\text{um}}) + \varepsilon_i,$$

the reduced residual $r_i$ generally still contains perturbation-induced variation through the difference

$$m_i(\boldsymbol{z}, \boldsymbol{D}) - \bar{m}_i(\boldsymbol{z}, D_i).$$

Therefore, unlike the structural-model result in Part I, identifiability through the reduced control function requires an additional condition ensuring that the latent nuisance contribution does not retain extra dependence on $\boldsymbol{D}$ after conditioning on $(\boldsymbol{z}, \boldsymbol{r})$.

**RC1 (Reduced first-stage identifiability).** For each $i$,

$$\bar{m}_i(\boldsymbol{z}, d) := \mathbb{E}[x_i \mid \boldsymbol{z}, D_i = d]$$

is identified from the joint distribution of $(x_i, \boldsymbol{z}, D_i)$.

**RC2 (Reduced-control-function bridge condition).** For each target node $j$, there exists a measurable function $\Lambda_j^{\mathrm{rcf}}(\boldsymbol{z}, \boldsymbol{r})$ such that

$$\mathbb{E}\big[\exp\big(\boldsymbol{z}^\top \boldsymbol{\alpha}_j + \phi_j(\boldsymbol{z}, \boldsymbol{z}_{\mathrm{um}})\big)\big|\boldsymbol{z}, \boldsymbol{D}, \boldsymbol{r}\big] = \Lambda_j^{\mathrm{rcf}}(\boldsymbol{z}, \boldsymbol{r}). \tag{20}$$

*Interpretation.* This is the key assumption specific to the reduced control function. It states that once $(\boldsymbol{z}, \boldsymbol{r})$ are given, the latent nuisance component is no longer affected by the perturbation pattern $\boldsymbol{D}$.

##### Reduced-control-function representation

**Lemma 2.** *Under RC1–RC2,*

$$\mathbb{E}[y_j \mid \boldsymbol{z}, \boldsymbol{D}, \boldsymbol{r}] = \exp\left(\sum_{k=1}^{p} \big(\bar{m}_k(\boldsymbol{z}, D_k) + r_k\big) w_{kj}^0 D_j^e + v_j D_j\right) \Lambda_j^{\mathrm{rcf}}(\boldsymbol{z}, \boldsymbol{r}). \tag{21}$$

*Interpretation.* This is the reduced-control-function analogue of (15). The perturbation dependence enters through the reduced first-stage means $\bar{m}_k(\boldsymbol{z}, D_k)$, while all latent nuisance terms are absorbed into $\Lambda_j^{\mathrm{rcf}}(\boldsymbol{z}, \boldsymbol{r})$.

*Proof.* By definition of $r_k$,

$$x_k = \bar{m}_k(\boldsymbol{z}, D_k) + r_k.$$

Substituting into (13) gives

$$\begin{aligned}\mathbb{E}[y_j \mid \boldsymbol{x}, \boldsymbol{z}, \boldsymbol{z}_{\mathrm{um}}, D_j] = \exp\Bigg(&\boldsymbol{z}^\top \boldsymbol{\alpha}_j + \phi_j(\boldsymbol{z}, \boldsymbol{z}_{\mathrm{um}}) \\ &+ \sum_{k=1}^{p} \big(\bar{m}_k(\boldsymbol{z}, D_k) + r_k\big) w_{kj}^0 D_j^e + v_j D_j\Bigg).\end{aligned} \tag{22}$$

Taking conditional expectation given $(\boldsymbol{z}, \boldsymbol{D}, \boldsymbol{r})$ and applying RC2 yields (21). □

##### Reduced-control-function identifiability under low-MOI

**RL (Reduced low-MOI relevance).** For each $i$,

$$\bar{\delta}_i(\boldsymbol{z}) := \bar{m}_i(\boldsymbol{z}, 1) - \bar{m}_i(\boldsymbol{z}, 0)$$

is nonzero on a set of positive probability.

**Theorem 3** (Reduced-control-function identifiability under low-MOI)**.** *Under RC1–RC2, ML1, and RL, the matrix $\boldsymbol{W}_0$ is identifiable under the low-MOI design.*

*Intuition.* This is the cleanest regime for the reduced control function. Comparing $\boldsymbol{D} = \boldsymbol{0}$ and $\boldsymbol{D} = \boldsymbol{e}_i$ with $i \neq j$ leaves both $\boldsymbol{r}$ and the nuisance factor $\Lambda_j^{\mathrm{rcf}}(\boldsymbol{z}, \boldsymbol{r})$ fixed, so the ratio isolates $w_{ij}^0$.

*Proof.* Fix $j$ and $i \neq j$. Under ML1, both $\boldsymbol{D} = \boldsymbol{0}$ and $\boldsymbol{D} = \boldsymbol{e}_i$ occur with positive probability. Since $i \neq j$, both states satisfy $D_j = 0$, hence $D_j^e = 1$. By (21),

$$\mathbb{E}[y_j \mid \boldsymbol{z}, \boldsymbol{r}, \boldsymbol{D} = \boldsymbol{0}] = \exp\left(\sum_{\ell=1}^{p} (\bar{m}_\ell(\boldsymbol{z}, 0) + r_\ell) w_{\ell j}^0\right) \Lambda_j^{\mathrm{rcf}}(\boldsymbol{z}, \boldsymbol{r}),$$

and

$$\mathbb{E}[y_j \mid \boldsymbol{z}, \boldsymbol{r}, \boldsymbol{D} = \boldsymbol{e}_i] = \exp\left((\bar{m}_i(\boldsymbol{z}, 1) + r_i) w_{ij}^0 + \sum_{\ell \neq i} (\bar{m}_\ell(\boldsymbol{z}, 0) + r_\ell) w_{\ell j}^0\right) \Lambda_j^{\mathrm{rcf}}(\boldsymbol{z}, \boldsymbol{r}).$$

Taking the log-ratio yields

$$\log \frac{\mathbb{E}[y_j \mid \boldsymbol{z}, \boldsymbol{r}, \boldsymbol{D} = \boldsymbol{e}_i]}{\mathbb{E}[y_j \mid \boldsymbol{z}, \boldsymbol{r}, \boldsymbol{D} = \boldsymbol{0}]} = \bar{\delta}_i(\boldsymbol{z})\, w_{ij}^0.$$

By RC1 and RL, $\bar{\delta}_i(\boldsymbol{z})$ is identified and nonzero on a set of positive probability, hence $w_{ij}^0$ is uniquely determined. Repeating this argument for all $i \neq j$ and all $j$ proves identifiability of $\boldsymbol{W}_0$. □

**Reduced-control-function identifiability under multi-perturbation designs**

For the multi-perturbation regime, the reduced-control-function identifiability argument again requires a coefficient-wise non-collinearity condition, now formulated in terms of the reduced first-stage means $\bar{m}_i(\boldsymbol{z}, D_i)$ rather than the full first-stage means $m_i(\boldsymbol{z}, \boldsymbol{D})$.

**RH1$_{ij}$ (Reduced coefficient-wise non-collinearity).** Fix $j$ and $i \neq j$. There exists $d_j \in \{0,1\}$ such that $D_j^e(d_j) = 1$ and

$$\mathbb{P}(D_j = d_j \mid \boldsymbol{z}) > 0$$

on a set of positive probability, and

$$\operatorname{Var}\Big(D_i - \mathbb{E}[D_i \mid \boldsymbol{z}, D_j = d_j, \boldsymbol{D}_{-(i,j)}] \;\Big|\; \boldsymbol{z}, D_j = d_j\Big) > 0 \tag{23}$$

on a set of positive probability.

**Theorem 4** (Reduced-control-function identifiability under multi-perturbation designs)**.** *Fix $j$. Under RC1–RC2 and RH1$_{ij}$ for every $i \neq j$, the $j$-th column of $\boldsymbol{W}_0$ is identifiable from the reduced-control-function model. If these conditions hold for every $j$, then $\boldsymbol{W}_0$ is identifiable.*

*Interpretation.* The reduced-control-function approach identifies the coefficient vector through variation in the perturbation indicators themselves, because the reduced first-stage means depend only on $D_i$. The condition RH1$_{ij}$ is coefficient-wise and allows some perturbation pairs to be incompatible.

*Proof.* Fix $j$. Suppose another vector $\tilde{\boldsymbol{w}}_{\cdot j}$ yields the same reduced-control-function model. From (21),

$$\sum_{k=1}^{p} (\bar{m}_k(\boldsymbol{z}, D_k) + r_k)(w_{kj}^0 - \tilde{w}_{kj}) D_j^e + (v_j - \tilde{v}_j) D_j + c_j(\boldsymbol{z}, \boldsymbol{r}) = 0$$

almost surely for some measurable $c_j$. Restrict to a stratum $\{D_j = d_j\}$ with $D_j^e(d_j) = 1$. Since $w_{jj}^0 = \tilde{w}_{jj} = 0$,

$$\sum_{k \neq j} (\bar{m}_k(\boldsymbol{z}, D_k) + r_k)(w_{kj}^0 - \tilde{w}_{kj}) + b_j(\boldsymbol{z}, \boldsymbol{r}, d_j) = 0.$$

Using

$$\bar{m}_k(\boldsymbol{z}, D_k) = \bar{m}_k(\boldsymbol{z}, 0) + \bar{\delta}_k(\boldsymbol{z}) D_k, \qquad \bar{\delta}_k(\boldsymbol{z}) := \bar{m}_k(\boldsymbol{z}, 1) - \bar{m}_k(\boldsymbol{z}, 0),$$

this becomes

$$\sum_{k \neq j} \bar{\delta}_k(\boldsymbol{z}) D_k (w_{kj}^0 - \tilde{w}_{kj}) + \tilde{b}_j(\boldsymbol{z}, \boldsymbol{r}, d_j) = 0.$$

Fix $i \neq j$, define

$$a_k(\boldsymbol{z}) := \bar{\delta}_k(\boldsymbol{z})(w_{kj}^0 - \tilde{w}_{kj}),$$

and project onto

$$\eta_i := D_i - \mathbb{E}[D_i \mid \boldsymbol{z}, D_j = d_j, \boldsymbol{D}_{-(i,j)}].$$

Multiplying by $\eta_i$ and taking conditional expectation given $(\boldsymbol{z}, D_j = d_j)$ gives

$$a_i(\boldsymbol{z})\, \mathbb{E}[\eta_i^2 \mid \boldsymbol{z}, D_j = d_j] = 0.$$

By RH1$_{ij}$,

$$\mathbb{E}[\eta_i^2 \mid \boldsymbol{z}, D_j = d_j] > 0$$

on a set of positive probability, hence $a_i(\boldsymbol{z}) = 0$. Therefore

$$\bar{\delta}_i(\boldsymbol{z})(w_{ij}^0 - \tilde{w}_{ij}) = 0.$$

If $\bar{\delta}_i(\boldsymbol{z}) \neq 0$ on a set of positive probability, then $w_{ij}^0 = \tilde{w}_{ij}$. Since $i \neq j$ was arbitrary, the $j$-th column is identifiable. □

**Justification of RC2 in the present setting.**

We now explain why the bridge condition RC2 is reasonable under the structural model considered in this work.

Recall that under the relaxed first-stage model,

$$x_i = m_i(\boldsymbol{z}, \boldsymbol{D}) + h_i(\boldsymbol{z}, \boldsymbol{z}_{\mathrm{um}}) + \varepsilon_i, \tag{24}$$

and the reduced control-function residual is defined as

$$r_i := x_i - \bar{m}_i(\boldsymbol{z}, D_i), \qquad \bar{m}_i(\boldsymbol{z}, D_i) := \mathbb{E}[x_i \mid \boldsymbol{z}, D_i].$$

Therefore,

$$r_i = \underbrace{m_i(\boldsymbol{z}, \boldsymbol{D}) - \bar{m}_i(\boldsymbol{z}, D_i)}_{\text{perturbation-induced component}} + h_i(\boldsymbol{z}, \boldsymbol{z}_{\mathrm{um}}) + \varepsilon_i. \tag{25}$$

The role of the reduced control function is to capture the component of $\boldsymbol{x}$ that is driven by latent confounding through $h_i(\boldsymbol{z}, \boldsymbol{z}_{\mathrm{um}})$. The key observation is that, under mild conditions, conditioning on $(\boldsymbol{z}, \boldsymbol{r})$ effectively controls for $\boldsymbol{z}_{\mathrm{um}}$ up to noise, while the remaining dependence of $\boldsymbol{r}$ on $\boldsymbol{D}$ does not convey additional information about $\boldsymbol{z}_{\mathrm{um}}$.

More precisely, RC2 requires that

$$\mathbb{E}\big[\exp\big(\boldsymbol{z}^\top \boldsymbol{\alpha}_j + \phi_j(\boldsymbol{z}, \boldsymbol{z}_{\mathrm{um}})\big) \mid \boldsymbol{z}, \boldsymbol{D}, \boldsymbol{r}\big] = \Lambda_j^{\mathrm{rcf}}(\boldsymbol{z}, \boldsymbol{r}), \tag{26}$$

i.e., that the conditional distribution of $\boldsymbol{z}_{\mathrm{um}}$ given $(\boldsymbol{z}, \boldsymbol{D}, \boldsymbol{r})$ depends on $(\boldsymbol{z}, \boldsymbol{r})$ but not further on $\boldsymbol{D}$.
